# Heads in the clouds: marine viruses disperse bidirectionally along the natural water cycle

**DOI:** 10.1101/2022.06.21.497027

**Authors:** Janina Rahlff, Sarah P. Esser, Julia Plewka, Mara Elena Heinrichs, André Soares, Claudio Scarchilli, Paolo Grigioni, Heike Wex, Helge-Ansgar Giebel, Alexander J. Probst

## Abstract

Marine viruses have thoroughly been studied in seawater, yet their dispersal from neuston ecosystems at the air-sea interface towards the atmosphere remains a knowledge gap. Here, we show that 6.2 % of the studied virus population were shared between air-sea interface ecosystems and rainwater. Virus enrichment in the 1-mm thin surface microlayer and sea foams happened selectively, and variant analysis proved virus transfer to aerosols and rain. Viruses detected in rain and aerosols showed a significantly higher percent G/C base content compared to marine viruses, and a genetically distinct rain virome supports that those viruses could inhabit higher air masses. CRISPR spacer matches of marine prokaryotes to “foreign” viruses from rainwater prove regular virus-host encounters at the air-sea interface. Our findings on aerosolization and long-range atmospheric dispersal implicate virus-mediated carbon turnover in remote areas, viral dispersal mechanisms relevant to human health, and involvement of viruses in atmospheric processes like ice-nucleation.

---

Marine viruses represent the most abundant biological entities in the oceanic water column ^1^ where they contribute to microbial diversity ^2^, supply hosts with metabolic genes ^3^ and influence carbon cycling by inducing host cell lysis (the “viral shunt”)^4^. While viruses have been studied in many marine habitats including the surface water ^5^ and deep-sea sediments ^6^, their presence at the air-sea interface, where microorganisms modulate gas and organic matter exchange processes ^7–9^, remains mostly enigmatic. Like many micro- and macroorganisms ^10,11^, viruses accumulate in the thin (<1 mm) uppermost layer of aquatic ecosystems, the surface microlayer (SML, reviewed by Cunliffe, et al. ^12^,Engel, et al. ^13^), where they belong to a pool of organisms collectively referred to as neuston ^14^. The enrichment of the virioneuston in the SML is mediated by bubble transport from the underlying water ^15,16^ and likely maintained by viral attachment to particles ^17^ and a dependency on abundant prokaryotic hosts ^18^. In freshwater, bacteriophages residing in the SML can form autochthonous communities ^19^ but comparatively little viromics studies have been conducted for marine SML (reviewed by Rahlff ^20^). Sea foams float as (extended) patches on the sea surface (Supplementary Video), forming deposits at the shoreline and being microbial habitats that contrast the SML ^21,22^. Based on satellite data, foams (whitecaps) cover up to 6 % of oceanic surface area and are expected to become more frequent with climate change ^23^. During storms, foams can flood beaches ^15^ and massively pollute coastal areas like recently in Turkey ^24^. Furthermore, sea foams can contain pathogenic bacteria ^25^ and their easy spread might be an important step for the dispersal and aerosolization of its inhabiting microbes ^26^ and potentially viruses. Foams can effectively concentrate viruses ^27^ which survive more than three hours of drying and sunlight when caught in foams ^28^. Virus-like particles (VLP) can reach a 300 x higher abundance in foams compared to surrounding waters ^24^.

Interest in studying viruses in the skin-like layer between ocean and atmosphere arises from therein appearing human pathogenic viruses ^29^, the potential of SML viruses to get airborne ^30^, to selectively enrich in aerosols ^31^, and to disperse over long distances to eventually promote turnover of algal blooms in remote regions ^32^. Virus aerosolization from the SML was previously studied ^15,30,33^, but investigations pursuing metagenomic approaches to explore the virioneuston and its aerosolization in the field are lacking. A recent review highlighted the need to quantify marine aerobiota, to characterize the spatial-temporal dimensions of dispersal, and to understand acclimations of marine microorganisms to atmospheric conditions ^34^. Once airborne, viruses could even fulfill other functions as recently suggested ^35^: Airborne marine viruses could serve as ice-nucleating particles (INP), a function already described for many microorganisms ^36,37^, and act as catalysts to mediate freezing at temperatures warmer than 10 °C. INP exist in the SML ^38–40^ and have an important role in cloud formation, cloud albedo and precipitation and thus are key in climate regulation dynamics ^41^. Viruses and bacteria can be found in clouds ^42–45^, where the latter might grow selectively ^46^ and as INPs, trigger their own precipitation ^47,48^. Precipitation could be an underestimated source of microorganisms to Earth’s surface, for example it contributed to as much as 95% of atmospheric bacterial deposition at a Korean site ^46^. So far, research on viruses included in wet precipitation was mainly focused on viruses relevant to human health, such as enteric or adenoviruses ^49–51^. Reche, et al. ^52^ reported that 10^7^ bacteria and 10^9^ viruses per m^2^ per day deposit above the atmospheric boundary layer, with marine origin making stronger contributions than terrestrial sources. This rate can be one order of magnitude higher for bacteria ^53^ and perhaps also for viruses. Rain events related to a hurricane decreased marine viral diversity and abundance as well as introduced new taxa in the western Gulf of Mexico ^54^. Furthermore, stormwater runoff changed the viral community composition of inland freshwaters and a stormwater retention pond ^55,56^. However, the transmission cycle of neustonic viruses from the sea surface via aerosols to wet precipitation remains speculative and the degree of viral exchange between these habitats and along the natural water cycle is unknown.

To fill these knowledge gaps, we analyzed 55 metagenomes including air-sea interface habitats (sea foams and SML), subsurface water (SSW) from 1 m depth, aerosols, and precipitation (rain, snow) collected in a coastal region of the Skagerrak in Tjärnö, Sweden (Fig. 1A). We explored the potential of marine viruses to become aerosolized and being returned to Earth via wet deposition. We hypothesized that aerosols and rain included marine viruses, and that sea foams have a particular role in the contribution to viral aerosolization due to their high content of organic material, high air content, and direct contact with the atmosphere. Virus-host infections histories were investigated by studying the prokaryotic adaptive immunity, namely the clustered regularly interspaced short palindromic repeats (CRISPR) system. Extraction and matches of CRISPR spacers, short sequences obtained from cell-invading mobile genetic elements, to viral protospacers allowed to reveal virus-host relationships across ecosystem boundaries.

**Figure 1:**
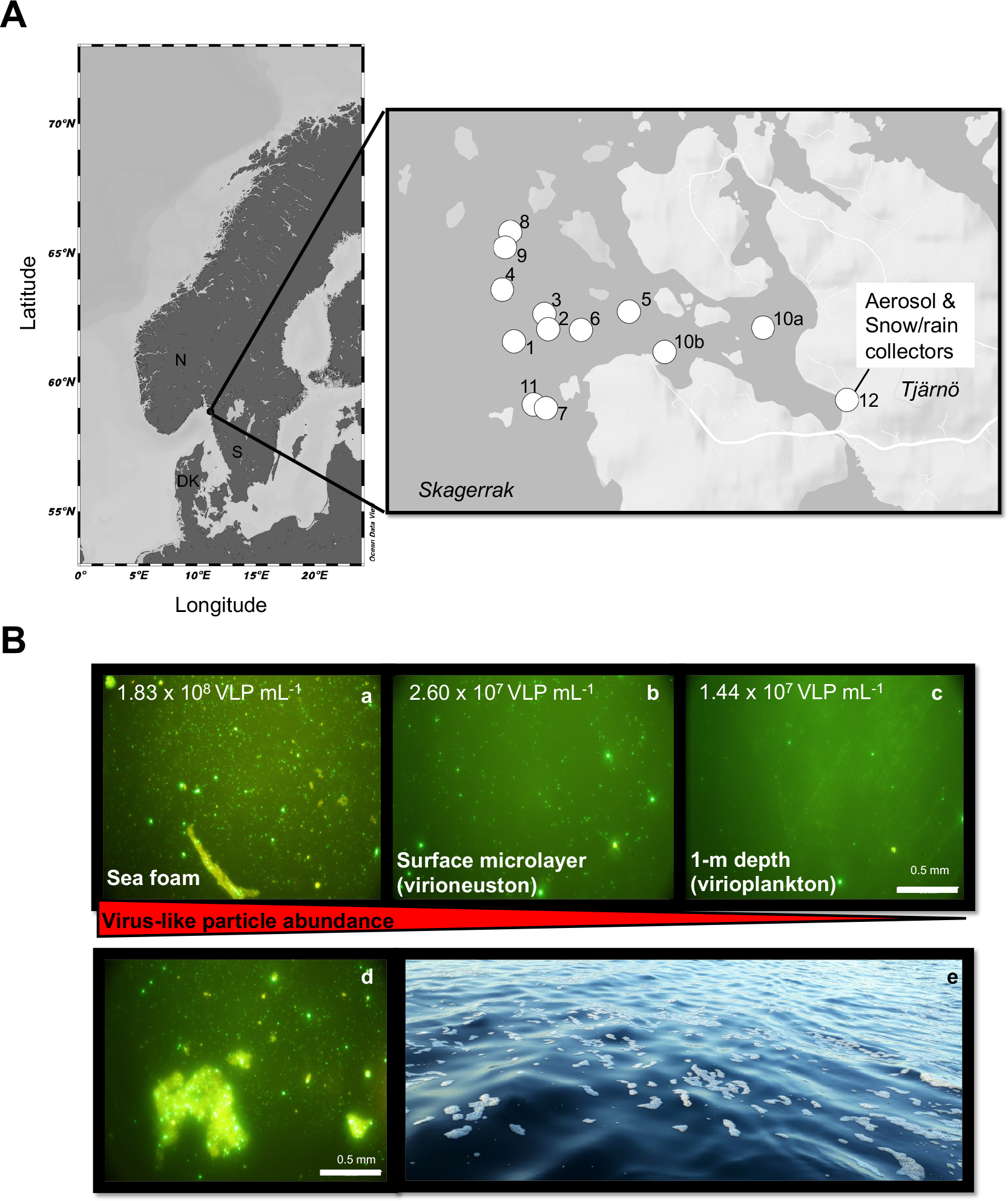
Map depicting sampling stations and viral enrichment in sea-surface microlayer and sea foam. Map of sampling sites was generated using Ocean Data View ^118^ and https://maps.co. For further details about the stations, please refer to Table S9 (A). Gradient of virus-like particles (VLP) in sea foam (a), sea-surface microlayer (b) and 1-m deep subsurface water (c) recorded in epifluorescence microscopy with VLP counts mL^−1^ from Station 4 as obtained from flow cytometry. Virus-like particles stick to particulate matter in the foam habitat (d). Sea foams were collected as floating patches from the ocean’s surface (e) (B).

## Results

### Cell and VLP abundance, enrichments, and correlations thereof reveal tight virus-host coupling in the neuston

Marine, aerosol, and rain samples were collected around Tjärnö Marine Laboratory, Sweden including 11 stations in coastal waters of the Skagerrak, where air-sea interface samples (SML, foam) and a reference depth were sampled (Fig. 1A). Prokaryotic, small phototrophic eukaryotic and VLP counts were measured to assess virus-host ratios and correlations in the neuston (SML, foam) compared to the underlying plankton in the SSW. Across all stations, flow cytometry revealed viral abundance between 5.0 × 10^7^ − 1.8 × 10^8^, 1.3 × 10^7^ − 3.4 × 10^7^ and 1.4 × 10^7^ − 2.0 × 10^7^ VLPs mL^−1^ in floating sea foams, the SML and SSW, respectively (Extended Data Table 1, Extended Data Fig. 1). This increasing VLP gradient towards the atmosphere was supported by microscopic analysis as shown for station 4 (Fig. 1B a-c, Fig. S1), and VLPs in sea foams often adhered to particulate matter (Fig. 1B d). Counts of VLP in precipitation samples (rainwater) ranged between 3.7 × 10^4^ − 3.4 × 10^5^ VLPs mL^−1^. Enrichment factors (EF) for VLPs in the SML over SSW varied between 0.7 (depletion) and 1.9 (enrichment) (Extended Data Table 1). Total cell numbers of prokaryotes were 1.3 × 10^6^ − 3.8 × 10^6^, 7.0 × 10^5^ − 1.1 × 10^6^ as well as 7.0 × 10^5^ − 8.7 × 10^5^ cells mL^−1^ in floating sea foams (Fig. 1B e), the SML and SSW, respectively (Extended Data Fig. 1, Extended Data Table 1). EFs for prokaryotes fluctuated between 0.9 and 1.3. Across the five precipitation samples, 2.3 × 10^3^ − 1.8 × 10^4^ prokaryotic cells mL^−1^ were detected. Virus-host ratios (host=prokaryotes) based on flow cytometry data were highest in foams (range = 25.3 − 48.4), followed by the SML (range = 15.5-34.2) and SSW (range = 19.3 − 26.7). Virus-host ratios in precipitation samples showed the strongest variation and ranged between 7.1 and 127.8 (Extended Data Table 1). The highest virus-host ratios in the SML were detected on days were VLP EFs were ≥ 1.8 and prokaryotic EFs ≥ 1.1 at the same time. Total cell numbers of small phototrophic eukaryotes ranged between 3.4 × 10^3^ − 1.8 × 10^4^, 1.8 × 10^3^ − 6.2 × 10^3^, 2.1 × 10^3^ − 5.0 × 10^3^ cells mL^−1^ in sea foams, SML and SSW, respectively.

Within the SML, the number of small phototrophic eukaryotes and prokaryotes was significantly correlated with VLP abundance (Pearson’s corr = 0.74, t = 3.13, *p* = 0.014, df = 8, n = 10 and Pearson’s corr = 0.70, t = 2.75, *p* = 0.025, df = 8, n = 10 respectively, Extended Data Fig. 2 A&B), while the plankton/SSW correlations with the same variables were not significant (Extended Data Table 2, Extended Data Fig. 2 C & D). In addition, absolute numbers of small phototrophic eukaryotes and their EFs were significantly correlated with absolute numbers and EF of prokaryotes for the neuston, inferring a common transfer mechanism of these cell types to the air-sea interface (Extended Data Table 2, Fig. S2). EFs of VLP were significantly correlated to EFs of prokaryotes (Spearman’s rho = 0.62, *p* = 0.006, n = 10) but not to EFs of small phototrophic eukaryotes (Extended Data Fig. 2 E & F) probably indicating that enrichments of viruses in the SML are primarily dependent on host cell availability and that most SML viruses are prokaryotic viruses.

### Correlations of cell and VLP counts with ice-nucleating particles, wind speed and environmental parameters

INPs, like VLPs, showed a decreasing gradient from sea foams, SML to SSW (Extended Data Fig. 3), with INP in foams showing the highest concentrations over the detectable temperature range. Ice activity for all samples generally started at high temperatures of ~ −4 to −6 °C, comparable to observations for microorganisms in the atmosphere ^57^. Concentrations of INPs were only significantly correlated with VLPs, small phototrophic eukaryotes and prokaryotes, when all interface samples (sea foams and SML) were combined (Extended Data Table 2, Extended Data Fig. 4), but were not significantly correlated with VLPs and cells in the SSW. In addition, enrichment of VLPs (but not cells) in the SML was significantly correlated with the enrichment of INPs (Spearman’s rho = 0.86, *p* = 0.011, n = 8 Extended Data Table 2). Correlation analysis of absolute VLP counts derived from neuston and plankton as well as EFs for VLPs and cells with meteorological data (light, salinity, and wind speed) did not reveal any significant relationships (Extended Data Table 2). However, numbers of small phototrophic eukaryotes (Pearson’s corr = 0.68, t = 2.59, *p* = 0.031, *df* = 8, n = 10) and prokaryotes (Pearson’s corr = 0.71, t = 2.83, *p* = 0.022, *df* = 8, n = 10) from the SSW but not the SML were significantly correlated with salinity, which could be explained by regular inflow of high saline water from the Atlantic Ocean that probably affects deeper water layers more than the SML.

We applied linear models to investigate combinatory effects of environmental variables (wind speed, light, and salinity) on enrichment of cells and VLPs in the SML. One linear model considering the combinatory effects of wind speed and salinity on the enrichment of small phototrophic eukaryotes in the SML passed the F-test with significance (F = 5.43, *p* = 0.038, *df* = 6), and 59.6 % of the residuals were explained by this model. The model’s Akaike information criterion (AIC) was −2.37, which was superior to considering wind speed (AIC = 5.84) and salinity (AIC = 6.40) alone. Other models testing single and combined environmental parameters on the enrichments of cells and VLP in the SML were not significant.

### Aerosolization of biota and decreasing diversity from marine towards atmospheric habitats

Analysis of Shannon-Wiener Diversity Index of prokaryotic communities (extractions from filters with 5 − 0.2 μm pore size) suggests a lower diversity of atmospheric (rain, aerosol) microbiota compared to marine samples, with the difference between aerosols and SSW being significant (Kruskal-Wallis with Dunn’s multiple comparisons test, *p* = 0.0097, Fig. 2A). Permutational multivariate analysis of variance (PERMANOVA) confirmed significant differences for the non-metric multidimensional scaling (NMDS) analysis (*p* < 0.001, Fig. 2B) with TukeyHSD on multivariate dispersion distances revealing significant differences between aerosols and foam (*p* = 0.02), aerosols and SML (*p* = 0.02), and aerosols and SSW (*p* = 0.0002) (Fig. S3). Correlation matrices showed that aerosol and rain communities are distinct from marine communities and from each other (Extended Data Fig. 5A). Beta-diversity analysis showed that SSW samples were mainly composed of Proteobacteria (mean relative abundance ± standard deviation = 67.8 ± 5.2 %, n = 9), Bacteroidetes (22.6 ± 6.0 %), and Thaumarchaeota (5.1 ± 2.3 %, Fig. 2C). In general, the SML samples reflected this composition although two samples deviated by containing a major percentage of Planctomycetes (13.3 ± 28.7 %, n = 9) and Cyanobacteria (4.6 ± 7.2 %, n=9). Sea foams were like the SML but additionally contained WOR-2 (2.7 ± 3.2 %, n=3) and an increasing proportion of Bacteroidetes (37.7 ± 11.6 %, n = 3). Aerosols also contained Proteobacteria (43.4 ± 13.9 %, n = 8), Bacteroidetes (21.0 ± 21.6 %) and Planctomycetes (20.8 ± 13.0 %). The snow sample contained a relative abundance of 97.3 % Cyanobacteria (*Rivularia* sp.). The mean relative abundance of Proteobacteria (49.9 ± 5.1 %) and Bacteroidetes (20.7 ± 2.4 %) in rain was comparable to aerosols, but in contrast to sea surface water and aerosols, Actinobacteria (6.3 ± 9.4 %) and Cyanobacteria (13.6 ± 5.1 %) were more abundant (Fig. 2C).

**Figure 2:**
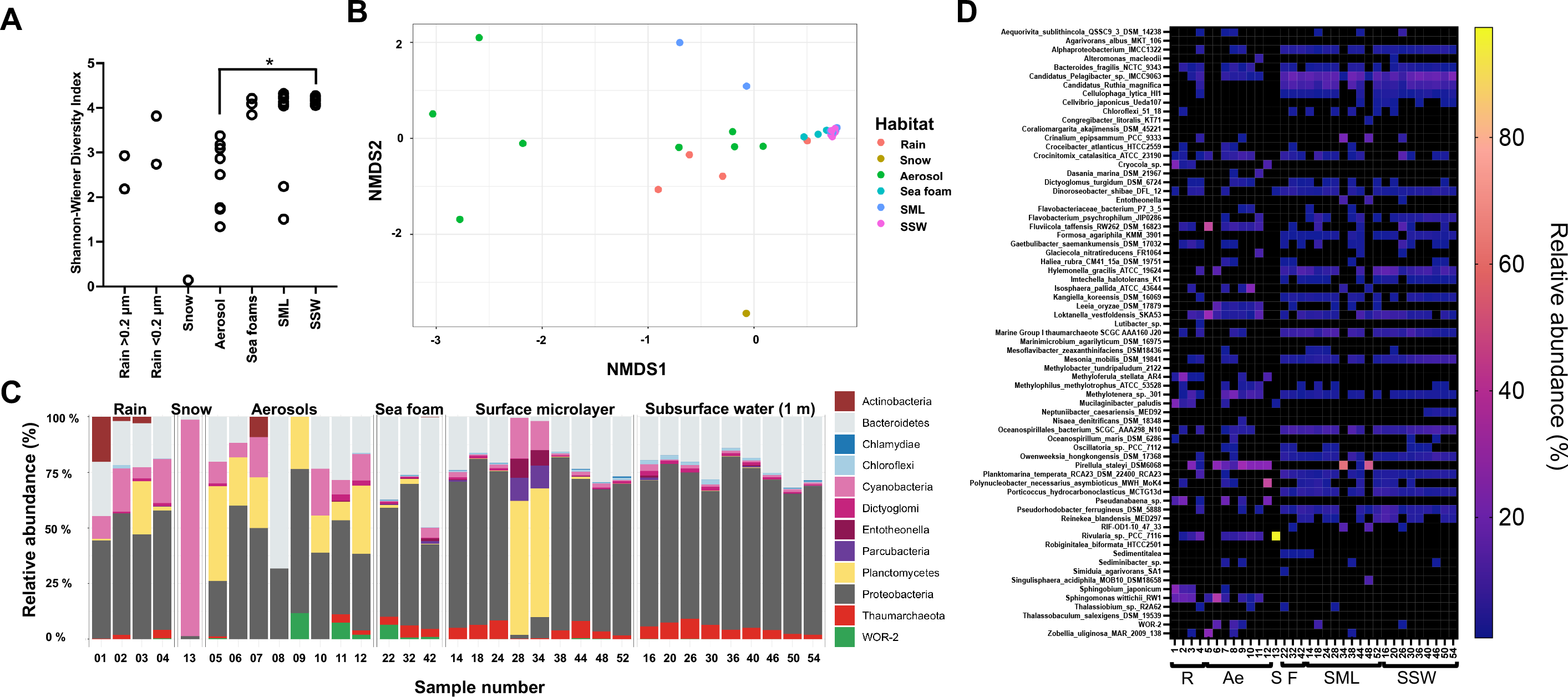
Diversity and relative abundance of marine and airborne prokaryotes based on relative abundance in rain, snow, aerosols, sea foam, surface microlayer (SML) and subsurface water (SSW). Diversity depicted by Shannon-Wiener index with * = p < 0.05 in Dunn’s multiple comparison test (post hoc analysis after Kruskal-Wallis test) (A), non-metric multidimensional scaling plot based on Bray-Curtis dissimilarity (stress = 0.082) (B), and stacked bar chart on beta-diversity at the phylum level (C). In C, relative abundance is based on read-normalized coverage on scaffolds carrying the ribosomal protein 3 gene (rps3) as explained in the main text. Black areas represent accumulated taxonomic units of minor abundance. Seawater samples show result of > 0.2μm samples, whereas rain contains >0.2 μm (#1+#3) and viromes (#2+#4). The heat map shows relative abundance of prokaryotic taxa across different ecosystems (D). Black colored fields indicate < 1% relative abundance; (R=rain, S=snow, Ae=aerosols, F=sea foams, SML=surface microlayer, SSW=subsurface water. Sample number is in accordance with Table S11;)

Detection of the same bacterial *ribosomal protein S3* genes in marine, aerosol and precipitation samples suggests their aerosolization from the sea surface, e.g., for Proteobacteria (*Oceanospirillum maris*, *Loktanella vestfoldensis, Candidatus* Pelagibacter), Bacteroidetes (*Crocinitomix catalasitica* and *Bacteroides fragilis*), Cyanobacteria (*Crinalium epipsammum* and *Oscillatoria* sp.) and Planctomycetes (*Pirellula staleyi*) (Fig. 2C&D). Interestingly, aerosols and precipitation contained Cyanobacteria such as *Rivularia* sp. (max. 14.1 % in rain, 97.7 % in snow) and Proteobacteria such as *Sphingobium japonicum* (max. 21.8 %) or *Methyloferula stellata* (max. 21.4 %), which could not be found in any of the local marine samples (relative abundance = 0). Rain contained Actinobacteria (*Cryocola* sp., max. 20.3 %) and Bacteroidetes such as *Mucilaginibacter paludis* (max. 20.1 %) that were only scarcely detected in marine samples (< 0.2 %) and thus probably originated from other sources.

### K-mer based virus-host assignments reveal *Pelagibacter* and *Porticoccus* as prevalent hosts

In total, 116 metagenome-assembled genomes (MAGs) could be recovered from 24 different samples (Table S1 & S2), which ranged from 58.8 – 100 % completeness (median = 86.3 %)and 0 – 11.8 % contamination (median = 3.9 %) based on uBin ^58^. CheckM ^59^ resulted in completeness and contamination scores of 18.4 – 99.5 % (median = 80.6 %) and 0 – 17.3 % (median = 1.6%) for these MAGs, respectively. Most host MAGs were of bacterial origin, except for three assigned to the genus *Nitrosopumilus* (Archaea). Recovering MAGs from atmospheric samples was particularly challenging with only a single MAG obtained from rain (genus *Pedobacter*) and from an aerosol sample (class *Planctomycetes*, order *Pirellulales*), respectively. Overall, bacterial MAGs were mostly classified as Gammaproteobacteria (n = 43), Alphaproteobacteria (n = 30), Bacteroidia (n = 36), and Planctomycetes (n = 4). Based on read mapping and breadth, all MAGs were detected in a marine ecosystem (except for the *Pedobacter* sp. MAG), rain and some additionally in aerosols. (Table S3). MAGs were matched to viruses based on shared k-mer frequency patterns, revealing that 120 marine viruses matched a MAG assigned to *Candidatus* Pelagibacter (Extended Data Fig. 6). Hosts of rain viruses (not detectable in other sampled ecosystems) and one aerosol virus were predicted as MAGs belonging to the family *Porticoccaceae* and a *Flavobacterium*.

### Viral diversity and transfer from the sea surface to aerosols and rain

Shannon-Wiener index values were significantly different for viruses between aerosols and SSW (Dunn’s multiple comparison test, *p* < 0.0001), but also weakly different between SML > 0.2 μm fraction and SSW virome samples (*p* = 0.029) (Fig. 3A). The distinct viral community of SML > 0.2 μm samples was also demonstrated by beta-diversity analysis (Fig. 3 B&C). Here, PERMANOVA confirmed significant differences for the NMDS analysis (*p* = 0.001), and *post hoc* pairwise analysis revealed significant differences between the SSW virome and the SML 0.2 μm fraction (*p* = 0.013, Fig. S3). We investigated further on SML and foam viral clusters (VC) that were detected in aerosols and rain. Rainwater contained the abundant cluster VC_723_0 absent in samples from the other ecosystems (Fig. 3C) and with one associated scaffold related to *Rhizobium* phage RHph_N3_2. EFs were overall higher for VCs in rain (EF max. = 15.8) compared to enrichments in aerosols in reference to SML and foam (EF max. = 2.8, Fig 3D). VC_880_0 (max. EF = 7.0), VC_738_0 (max. EF = 7.8), and VC_771_0 (max EF = 5.4) were strongly enriched (coverage ratio > 1) in rain compared to foam and/or SML but contained only viruses from this study and were unrelated to viruses from public databases. Enriched in rain, overlap cluster (VC_634/VC_747) and overlap cluster (VC_746/VC_747) were both related to *Pelagibacter* phage HTVC023P (max. EF = 7.0, Table S4), whereas overlap cluster (VC_773/VC_829/VC_885) was related to *Flavobacterium* phage vB_FspM_immuto_3-5A (max. EF = 2.6). An overlap cluster refers to genomes sharing genetic overlap with other genome(s) belonging to multiple VCs. VConTACT2 detected various singletons and outliers, which usually represent new viruses, and had an EF > 8 for rain over sea surface ecosystems but were unrelated to any known virus. In aerosols, e.g., VC_970_0 (max EF = 1.2), VC_914_0 (max EF = 2.1), and VC_738_0 (max. EF = 1.9) showed slight enrichments compared to marine samples but were also unrelated to any known viruses (Fig. 3D). Three singletons and nine outliers were additionally enriched in aerosols with outliers (EF = 2.8 and 1.4) being associated with a virus related to *Methylophilales* phage Melnitz-1 EXVC043M and *Vibrio* phage vB_VorS-PVo5, respectively. Overall, assembled marine viruses shared protein clusters with *Synechococcus, Rhizobium*, *Cellulophaga, Flavobacteria, Vibrio* and *Pelagibacter* phages in vConTACT2^60^ (Table S4, Extended Data Fig. 7). The correlation matrix across all viromes and samples shows most positive correlations between foam, SML, SSW, which are well-interconnected systems, and some positive correlations of specific marine samples with aerosol and rain samples (Extended Data Fig. 5B). Viromes of marine samples and rain samples were sometimes even negatively correlated, suggesting alternative sources of rain viruses other than the sea surface.

**Figure 3:**
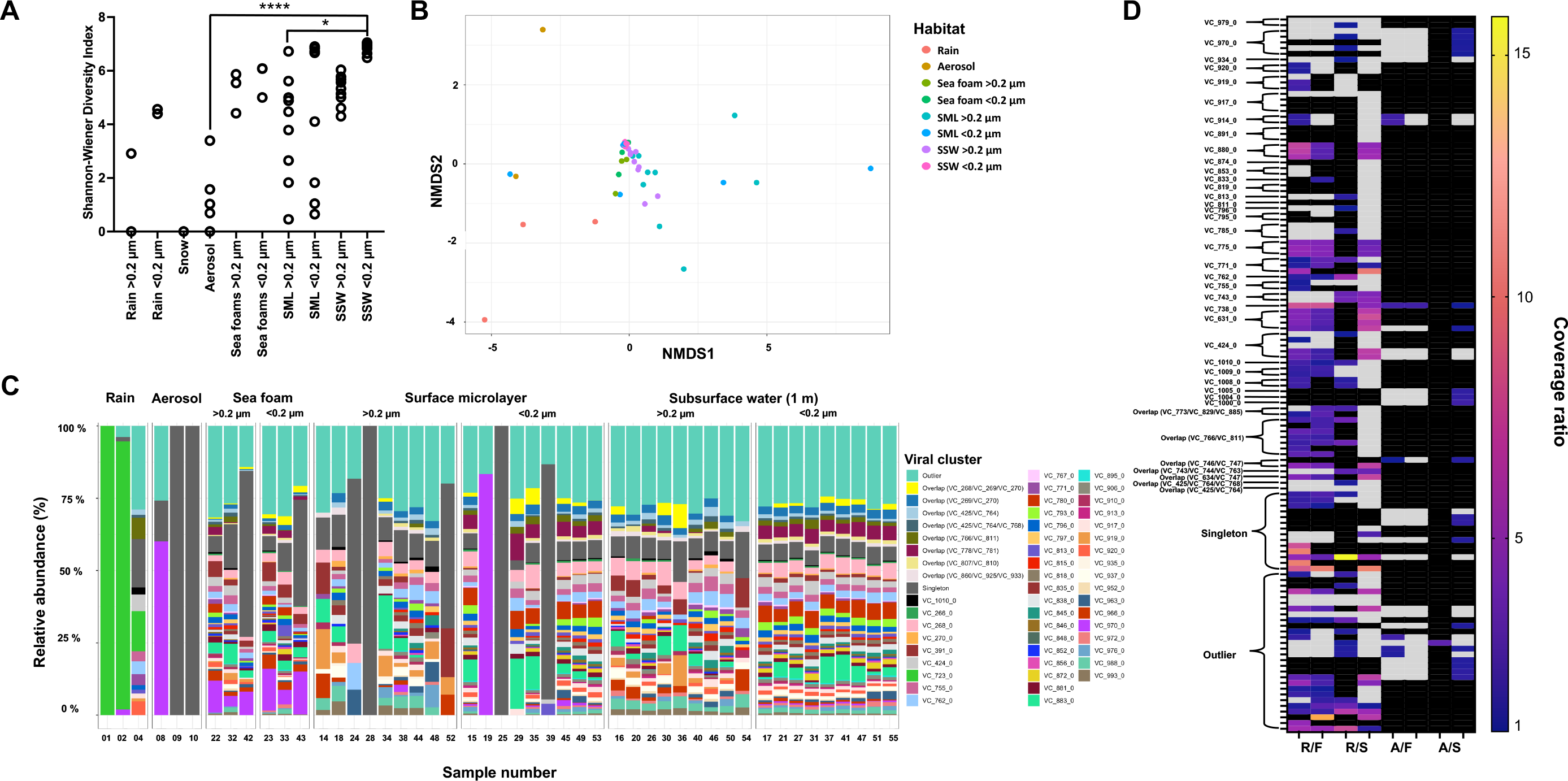
Diversity and enrichment analysis of marine and airborne viruses based on relative abundance in rain, snow, aerosols, sea foam, surface microlayer (SML) and subsurface water (SSW). Alpha-diversity for all samples (except for #12, which contains 0 viruses) depicted by Shannon-Wiener index; * = p < 0.05, **** = p < 0.0001 in Dunn’s multiple comparison test (post hoc analysis after Kruskal-Wallis test) (A), non-metric multidimensional scaling (NMDS) plot based on Bray-Curtis dissimilarity (stress = 0.06) (B), and stacked bar chart on beta-diversity (C). If samples only contained rare viruses (sample #7), a single virus (#3, #5, #6, #11 #13) or no viruses (#12), they were removed, and only the relative abundance of the 200 most abundant viruses assigned to viral clusters (VCs), outliers, and singletons were considered for B&C. In vConTACT2, outliers and singletons typically represent new viruses. In (C), marine samples are separated by size fraction: > 0.2 μm = prokaryote fractions, < 0.2 μm = viromes; Rain sample #1 is a > 0.2 μm sample, whereas #2 and #4 are rain viromes. Enrichment ratio of SML and foam viruses in rain and aerosols (D). Shown are ratios ≥ 1 of virus coverage for rain/foam (R/F), rain/SML (R/S), aerosol/foam (A/F), aerosol/SML (A/S), where the left tick stands for foam and SML virome and the right tick for foam and SML 0.2 μm fraction in the denominator. Black fields mean that the virus was absent in one or both ecosystems in the respective sample. Grey areas show out of range fields (ratio between 0 and 1, indicating depletion). Sample number is explained in Table S11.

We then investigated the aerosolization patterns of two circular viral genomes of similar length and carrying viral hallmark genes (terminases, portal protein) across the different habitats and stations (Fig. 4A+B). Virus_1 (39.7 kb, percent G/C content = 46.7 %, no RefSeq match in vConTACT2) was constantly of lower abundance in seawater samples compared to Virus_2 (35.1 kb, percent G/C content = 35.1 %, no RefSeq match in vConTACT2) across different stations. However, Virus_1 was consistently abundant in sea foams and was additionally found in three aerosol samples and two rain samples. Instead, Virus_2 was absent from the atmosphere despite its abundance in surface water. Virus_1 was linked to a *Porticoccus* MAG based on k-mer patterns (Extended Data Fig. 6) and single-nucleotide polymorphism (SNP) analysis revealed multiple SNP overlaps for this virus between a foam, aerosol, and rain sample, supporting its transfer from the sea surface to aerosols and rainwater (Fig. 4C&D, Table S5). Cross-mapping of reads from all samples against all 1813 viral scaffolds revealed shared viral populations between ecosystems (Fig. 6A). Sea foams, SML and SSW shared 837 viruses, whereas 15 viruses were present in all studied ecosystems. Overlaps between aerosols and rain samples must be treated with caution, because some rain could have reached the aerosol filter membrane during sampling and filter exchange (Table S6), although we tried to rule out the second possibility by subtracting reads from handling controls. Precipitation had the highest number of unique viruses (109), followed by foams (25), SSW (18), SML (7) and aerosols (6). Uniqueness means that no other habitat had 90 % identical reads with 75 % scaffold coverage of at least 1 × for that virus. A percentage of 6.2 % (111) of all viruses was shared between seawater including foam and precipitation. Interestingly, the rain sample pooled from Feb. 14^th^ to 22^nd^ of Feb. 2020 (event 2) defined most of this overlap compared to a sample from Feb. 7^th^ and 9^th^ February (event 1): Based on read-mapping, rain events 1 and 2 were associated with 22 versus 85 viruses assembled from marine samples as well as 112 versus 44 viruses assembled from rain, respectively. Based on read-mapping, event 1 delivered 38 marine prokaryotic MAGs (min. 90 % genome covered with reads), whereas 79 marine MAGs were found in the rain sample belonging to event 2 (Fig. 5). To explain these differences by tracking to potential sources, backward trajectories (TJs) for air masses were calculated. They showed that during event 2, air masses spent, during the first four days before arriving at the site, on average 72 % of their time over the sea and loading conditions (loading of air masses with generic particles) were fulfilled on average 35 % of the TJ. On the other hand, for event 1, air masses spent less time above the sea (64 %) and loading conditions were fulfilled, on average, only 10 % of the TJ points (Fig. 5).

**Figure 4:**
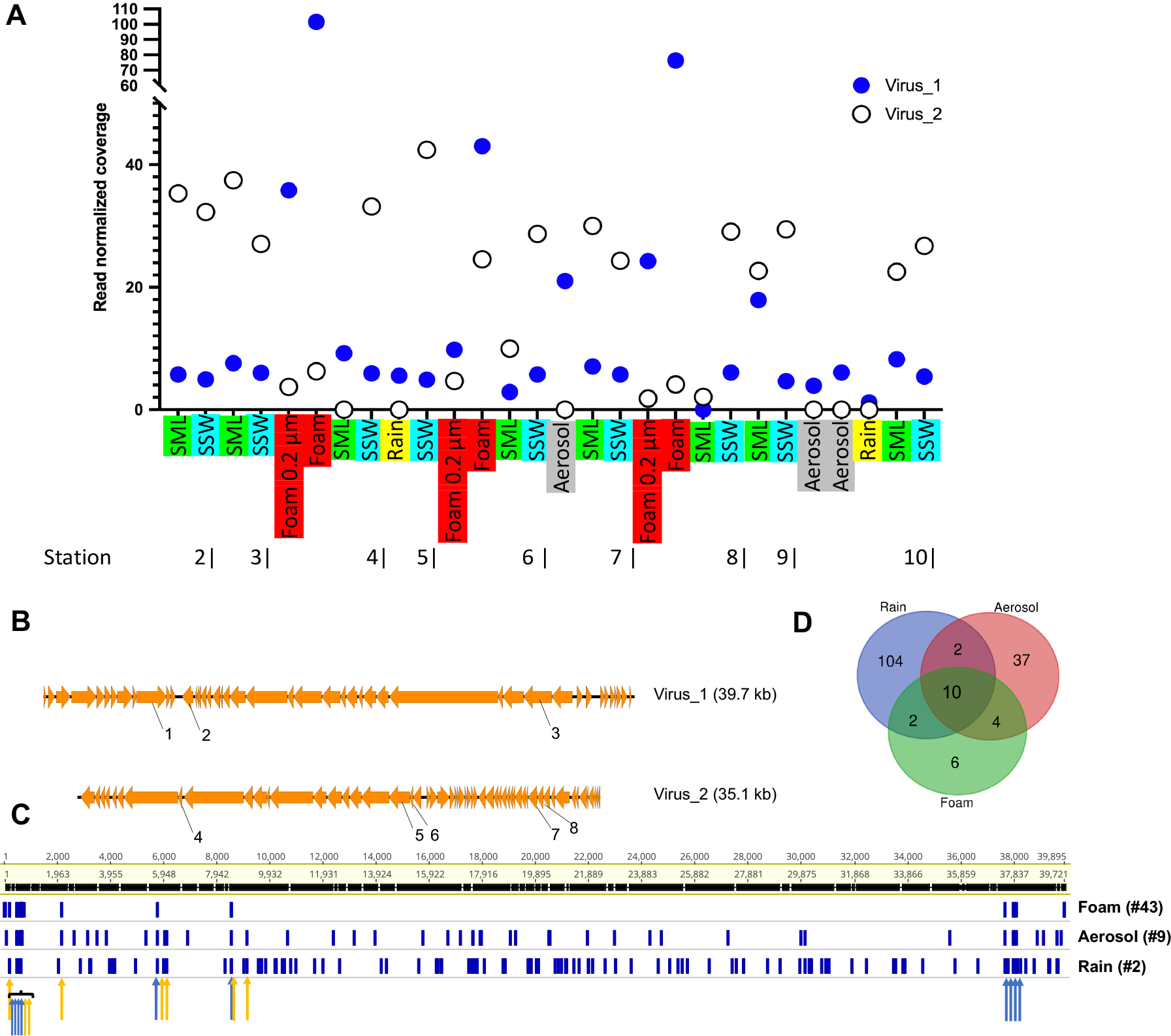
Succession of the coverage of two circular viral genomes across 28 metagenomes derived viromes of surface microlayer (SML), 1-m deep subsurface water (SSW) and sea foam as well as from sea foam filtered onto 0.2 μm membranes, aerosol, and rainwater samples (A). Synteny and functional annotations of the two circular viruses visualized using Easyfig ^82^ and annotated with DRAM-v^80^. Functional annotations are 1: Bifunctional DNA primase/polymerase, N-terminal [PF09250.12], 2: Sec-independent protein translocase protein (TatC) [PF00902.19], 3: Terminase, 4: Concanavalin A-like lectin/glucanases superfamily [PF13385.7], 5: Phage P22-like portal protein [PF16510.6], 6: Terminase-like family [PF03237.16]; Terminase RNAseH like domain [PF17288.3], 7: C-5 cytosine-specific DNA methylase [PF00145.18], 8: PD-(D/E)XK endonuclease [PF11645.9] (B). Variant analysis of Virus_1 for a sea foam, aerosol and rain sample reveals overlapping nucleotide polymorphisms. Blue and orange arrows indicate overlaps between three and two samples, respectively. For details, please see Table S4 (C). Venn diagram showing variant overlaps for Virus_1 in different ecosystems as shown in C (D).

**Figure 5:**
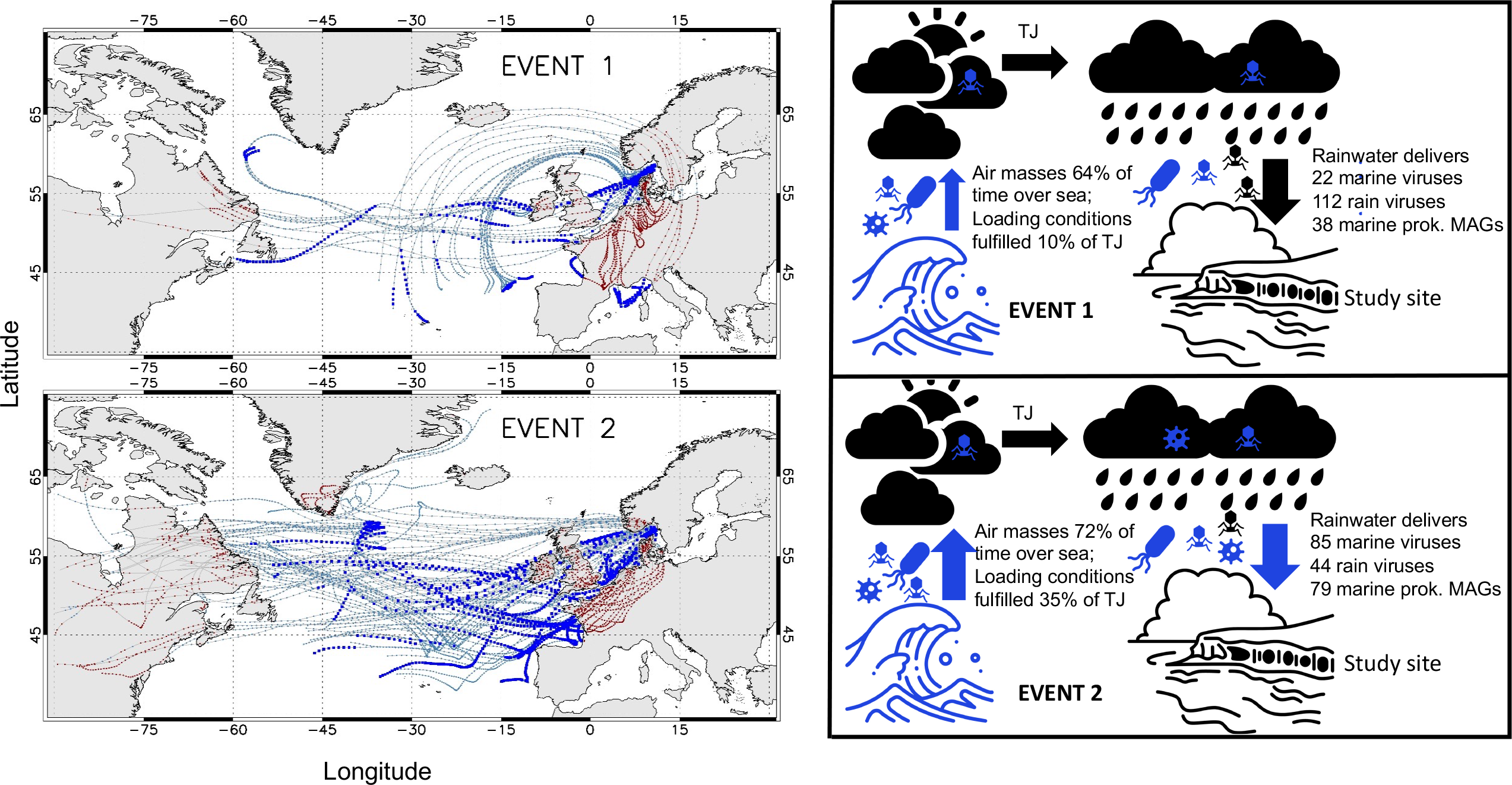
Backward trajectories (TJs) of two rain events leading to different deliveries of marine viruses and metagenome-assembled genomes (MAGs) at the study site. Event 1 (upper left panel) refers to a rainwater sample from 7^th^ to 9^th^ of Feb., and event 2 (lower left panel) to a sample collected between 14^th^ to 22^nd^ of Feb. 2020. Sky blue and red points highlight where backward trajectories travel above sea and land, respectively. Blue filled squares represent points where loading conditions were fulfilled (please see main text for further explanation). Panels on the right show corresponding deliveries of viruses and MAGs to the study sites. A virus was considered marine or from rain if assembled in such a sample and counted if detected based on read mapping. Marine MAGs were considered present in rainwater 0.2 μm samples if 90% of the genome was covered with reads (see Table S3).

**Figure 6:**
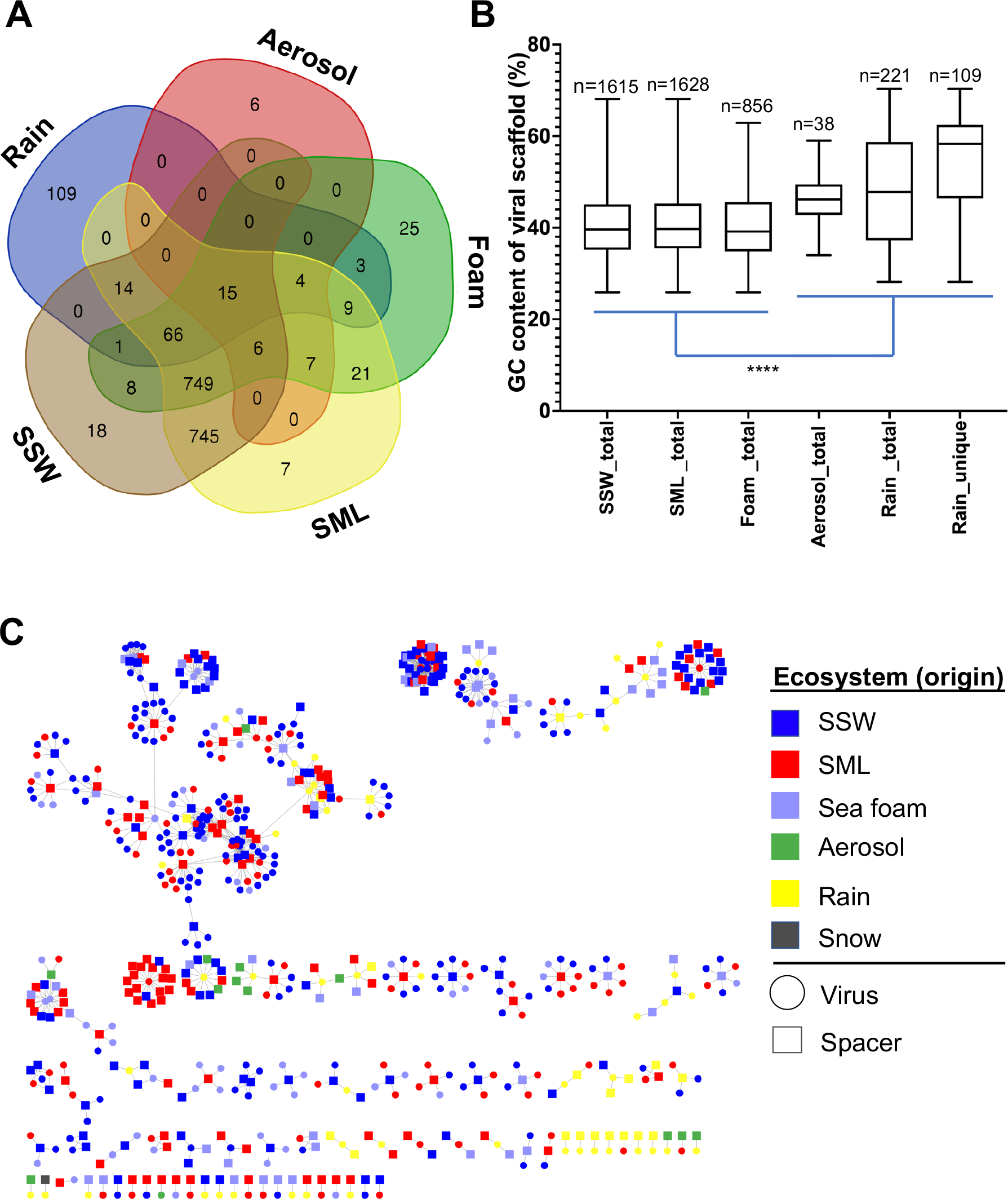
Overlapping occurrence of viral scaffolds, their percent G/C base content and CRISPR spacer to viral protospacer hits. Overview of shared viral scaffolds (>10 kb) length between seawater, aerosol and precipitation habitats obtained from 55 metagenomes and determined by mapping of reads. A viral genome was considered present in a sample if at least 75 % of the genome were covered with reads at least 90 % identical to the genome, in accordance with suggested viromics benchmarks ^78^ (A). Percent of the bases guanine (G) and cytosine (C) in viral scaffolds from rain, aerosol, foam, surface microlayer (SML) and subsurface 1-m deep water (SSW) based on read mapping. “Rain_unique” refers to viral genomes exclusively found in rain. Stars indicate significant differences after Kruskal Wallis test and Dunn’s multiple comparison test (****, p = <0.0001). In each pairwise comparison, the marine groups were significantly different from the atmospheric groups. Rain_total was also significantly different from Rain_unique (***, p = <0.001), which is not indicated to reduce complexity of the figure (B). CRISPR spacers (origin indicated as square) matching assembled viral scaffolds (circles) derived from different habitats (C).

### Rain and aerosol viruses show adaptations toward atmospheric residence and are targeted by marine prokaryote adaptive immunity

To investigate if unique atmospheric viruses have genetic adaptations, we explored the content of guanine (G) and cytosine (C) bases in viral scaffolds. Viral scaffolds solely occurring in rain samples (Rain_unique, n = 109) exhibited a significantly higher percent G/C content than total viruses found in rain (Kruskal-Wallis with Dunn’s multiple comparisons test, *p* = 0.0002). All viruses found in marine samples had a highly significantly lower percent G/C content compared to aerosol, total rain and unique rain viruses when compared pairwise (KW-test, *p* < 0.0001, Fig. 6B).

One very abundant circular viral genome was identified as a unique rain virus (VC_723_0, 39 kb, coverage = 189 x, percent G/C content = 59.7 %) with the closest relative of the VC being the *Rhizobium* phage RHph_N3_2. This phage carried typical phage hallmark genes like a major capsid protein, an endonuclease, a terminase and modification methylase, but also carried many hypothetical proteins (Table S7). In addition, two large viral scaffolds unique to rain (270 kb and 496 kb) and with some genes related to *Mimiviridae* and k-mer linked to a *Flavobacteriaceae* MAG, encoded for sensors of blue-light using FAD (BLUF, PFAM ID PF04940.13), a photoreceptor and for an UV-endonuclease UvdE (PF03851.15). Another 16 kb viral genome with typical phage proteins (terminase, capsid) encoded for Tellurium resistance genes TerD (PF02342.19). From metagenomic assemblies, and from one MAG of *Schleiferiaceae* bacterium MAG-54, CRISPR arrays with evidence level 3 and 4 from 18 different samples could be detected by CRISPRCasFinder ^61^, and mostly belonged to marine ecosystems (n = 14) and rainwater (n = 4, Table S8). CRISPR spacers extracted across all samples based on consensus direct repeat (DR) sequences from recovered arrays matched protospacers of viruses from seawater, but also the unique rain viruses (Fig. 6C, Fig. S4). Interestingly, CRISPR spacers matching most viral protospacers were extracted from two dominant arrays, with one of them targeting primarily (unique) rain viruses and the other one marine viruses (Fig S4, Table S8).

## Discussion

The atmosphere is known to mediate rapid transport of prokaryotes and viruses from marine and terrestrial sources over thousands of kilometers ^62,63^. Aerosol and rain samples contained MAGs and genetic signatures from marine microbes, suggesting that aerosolization from the sea surface generally took place. We found that precipitation samples for instance contained cyanobacteria of *Rivularia* sp. (order *Nostocales*), but this species could not be found in marine samples from the study site inferring transport from remote areas. Especially the rainwater community related to event 2 contained a mixture of local and probably dispersed prokaryotes. This confirms their travel along the natural water cycle and that the origin of the air mass is crucial for understanding airborne microbial diversity, especially near the ocean-atmosphere interface ^64^.

Due to the tighter coupling of VLPs to their hosts in the neuston, one of the determining factors for a virus to get aloft is probably being associated with a host cell or other particulates, such as transparent exopolymer particles. These are prone to aerosolization, were found in cloud water and to absorb viruses ^65,66^. Viral attachment to biotic and abiotic surfaces in seawater is a common phenomenon ^67^ and more likely to occur in the SML ^17^ and in foams (Fig. 1B). We found that VLPs in foams often adhered to particulate matter though >50 μm particles were removed prior flow cytometric measurements, probably leading to an underestimation of the measured VLP counts in particle-rich sea foams. Michaud, et al. ^33^ used a mesocosm experiment to show that bacterial and viral aerosolization is taxon-specific, and our data support and extend the concept of selective aerosolization from the water surface under field conditions. Two viral genomes experienced the same environmental conditions but showed a consistently distinct aerosolization behavior on a spatial-temporal scale. Virus_1 became constantly more abundant in sea foams at different stations, and SNP analysis proved its transfer to rainwater. As enrichments of viruses in rainwater over foam compared to rainwater over SML were enhanced in the > 0.2 μm fraction (Fig.3 D), aerosolization from sea foams as a “springboard” from sea to air represents a likely scenario. In addition, correlation analyses demonstrated independence of VLP enrichment in the SML from various environmental conditions, but findings only apply for wind speeds < 6 m s^−1^, since higher wind speeds prevent SML sampling and bacterioneuston enrichment ^68^. The impact of wind-induced sea spray formation on virus and host aerosolization, needs future investigations for instance by carrying out experiments in wind-wave facilities.

Atmospheric dispersal of viruses allows the spread of “foreign” genetic material into new habitats enabling bacterial evolution and explains why similar viral genomes can be found across large geographical distances ^69^. The fact that rain samples shared up to 6.2 % of the virome with marine samples, generally indicates profound viral exchange between both ecosystems. This is further supported by CRISPR spacers from sea surface prokaryotes matching viruses found exclusively in rain samples. Established adaptive immunity indicates a long-lasting virus-host relationship and proves a constant genetic inflow to the sea surface by precipitation.

Previous work reported that bacterial isolates from rain are often pigmented ^70^, aiding their survival at high UV radiation. Rain and aerosol viruses detected in this study had a significantly higher percent G/C base content compared to marine viruses, which is a suitable adaptation for avoidance of thymine-specific damage by UV radiation in bacteria^71^ and occurred in recently described bacterial isolates from the stratosphere ^72^. A viral cluster specific to rain was identified, and many viruses enriched in the atmosphere compared to seawater lacked representatives in public databases. Some viruses carried genes, e.g. for Tellurium resistance, which could be relevant for tolerating atmospheric conditions where Tellurium contaminations occur ^73^. As genetic adaptations cannot be established within hours, for instance shortly after aerosolization, we assume that the atmosphere contains its own virome, likely maintained across large areas and different altitudes, supplied by marine or terrestrial sources, and then distributed up to the higher troposphere or even above. The hereby detected 109 distinct rainwater viruses associated with rain event 1 and carrying genes related to atmospheric life supports this conclusion. In contrast, air masses delivering rain during event 2 were loaded over sea and contained more marine viruses and prokaryotic MAGs. We speculate that these viruses endure less well in the atmosphere than the high percent G/C-adapted rain viruses but can still be dispersed with air masses or clouds and deposited from them. Consequently, air mass and cloud migration and therein transported microbes and viruses could have strong impacts on biogeography, diversity, and evolution of microbial communities on the ground, particularly in aquatic habitats. Future work using culture-dependent experiments is required to elucidate if marine viruses remain infective when they are returned to the surface during precipitation. Since sea surface hosts possess CRISPR spacers matching genomes of unique rain viruses, ongoing infections with these viruses in the sea surface are probable. Moreover, there is a tendency that at ecosystem boundaries, different CRISPR arrays are in charge for targeting viruses from different origins (air, sea), although we assume that these arrays have high turnover rates in this dynamic interface ecosystem. Future work is certainly required to confirm the here reported trends. In conclusion, our study shows broad dispersal of microbes and viruses along the natural water cycle and emphasizes that viruses crossing interfaces and ecosystem boundaries, e.g., with rainwater, can have a crucial role for shaping Earth’s aquatic microbiomes.

## Online methods

### Seawater sampling and processing

Seawater sampling sites were located in the bay offshore Tjärnö, Swedish west coast in the Skagerrak (Fig.1), an area characterized by strong salinity gradients ^74^ (Table S9). Foams and SML were sampled from a small boat using the glass plate method ^75^ as previously described ^22^. Corresponding subsurface water (SSW) from 1 m depth was collected as a reference using a syringe connected to a weighted hose. Wind speed was measured with a handheld VOLTCRAFT AN-10 anemometer (Conrad Electronic, Hirschau, Germany), and light conditions were recorded on the boat using the Galaxy Sensors smartphone application v.1.8.10. Temperature and salinity were measured from the small boat using a portable thermosalinometer (WTW™ MultiLine™ 3420).

Water samples were stored in the dark and on ice until processing in the laboratory. Filtration equipment was treated prior to all usages with household bleach and rinsed with MilliQ water. Seawater (500 mL SML and 2 L SSW) and sea foams (200-400 mL) were sequentially vacuum filtered through 5 μm and 0.2 μm pore size Omnipore PTFE filter membranes (47 mm diameter, Merck/Sigma Aldrich, Munich, Germany). The flow-through of the 0.2 μm filter membrane was precipitated with 1 mg L^−1^ iron-III-chloride (Alfa Aesar/Thermo Fisher Scientific, Uppsala, Sweden) for 1 hour at room temperature ^76^, and the flocculates were in turn filtered onto another 0.2 μm Omnipore PTFE filter membrane to obtain viruses and small prokaryotes. All filters were stored at −80° C until further processing and shipped on dry ice to the home laboratory for DNA extraction from the 0.2 μm filter and the FeCl_3_ flocculates.

### Aerosol and precipitation sampling

We used a land-based aerosol pump/constant flow sampler (QB1, Dadolab, Milan, Italy) with a custom-made filtration unit (SIMA-tec GmbH, Schwalmtal, Germany) to filter aerosols from the atmosphere in coastal proximity (Fig. S5). Incoming air was filtered through 0.1 μm pore sized Omnipore PTFE filter membranes (Merck/Sigma Aldrich, Munich, Germany). Filtered volumes and filtration duration varied and ranged from 19 to 61 m^3^ (average volume flow 7 L min^−1^) and from 24 to 96.5 hours, respectively (Table S5). Gas volume was normalized to mean temperature and mean air pressure from start and end of an aerosol filtration. Handling controls for aerosol samples were collected as follows: a filter membrane was briefly placed on the filter unit, and directly frozen in a falcon tube at −80 °C. Snow and rain with a volume of 90 mL and 150 to 1050 mL, respectively were collected using funnels taped to Duran glass bottles. Rain was collected and, like seawater, filtered onto 0.2 μm pore size PTFE filter membranes, and the viral fraction was obtained as explained above. Rain collected between 14^th^ to 22^nd^ of Feb. 2020 was prefiltered onto 5 μm due to visible pieces (probably plant-based) in the sample. For enough DNA yield for sequencing, DNA from rain for the periods 07-09. Feb and 14^th^ to 22^nd^ of Feb. 2020 were pooled, respectively. The snow sample was prefiltered on 5 μm and frozen at −80 °C. Later in the home laboratory, it was thawed at room temperature and concentrated in an Amicon® Ultra-15 centrifugal filter unit (Ultracel 100 kDa) by spinning in several steps at 3000 x g, 10 minutes at 4 °C before DNA extraction.

### Air mass paths (backward trajectories)

Transport pathways of air masses was evaluated with 5-day backward trajectories (TJs) generated using the Hybrid Single-Particle Lagrangian Integrated Trajectories (HYSPLIT) model ^77^. The TJs were calculated every one hour ending at 700 m above the site for the period 1st to 29th February 2020. The European Centre for Medium-range Weather Forecasts (ECMWF) ERA5 model atmospheric reanalysis ^78^ is used to initialized HYSPLIT. After five days, the uncertainty associated with trajectories is estimated between 10 and 30 % of the travel distance^79^. Each TJ was then projected on the 10-m wind, total precipitation, land mask, surface pressure and cloud fraction model fields (ERA5), associating each point along the path with the nearest values of the considered model variables. The choice of the ending height (700 m) above the site is based on the analysis of *in situ* meteorological (Nordkoster A Automatic Weather Station, 58.890 °N, 11.010 °E as obtained from SMHI, https://www.smhi.se/en) and model (ERA5) data for February 2020 (Fig. S6). To identify loading areas and air masses presumably responsible for the transport towards the site, a selection of TJs was carried out considering those (ones) arriving above the site during precipitation sampling events 1 (7-9 February 2020) and 2 (14-16 and 20-22 February 2020). Similar to Becagli, et al. ^80^, loading conditions along TJs were evaluated searching where each TJ was within the mixing layer and wind speed at surface was greater than 3 m s^−1^.

### Microbial cell counts and virus-like particle abundances

Duplicates of unfiltered seawater, foam and precipitation samples were fixed with glutardialdehyde (1% final concentration, Merck, Sweden), stored for 1 hour in the dark and subsequently stored at −80 °C. Particle-enriched foams were gravity filtered onto 50 μm filters (CellTrics®, Sysmex Partec, Muenster, Germany) before cell counts of prokaryotes and small phototrophic (autofluorescent) eukaryotes were measured by a flow cytometer (BD Accuri C6, Becton Dickinson Biosciences, Franklin Lakes, USA) according to established protocols ^81–83^. Due to previously reported low coefficient of variance among SML biological replicates in flow cytometry ^68^, we did not measure biological replicates. VLPs were measured as done previously ^84^. The gating strategy is shown in Fig. S7. Flow cytometry results were further compared to VLPs counted under the epifluorescence microscope (Leica DMRBE Trinocular, Leica Microsystems). Enrichment factors (EFs) were calculated as previously performed ^22^. EF > 1 and < 1 indicate an enrichment and a depletion of measured specimens, respectively.

### Ice-nucleating particles

INP were measured from the 5 μm filter membrane that was used for pre-filtration of seawater samples. Of these filters, small disks with 1 mm diameter were punched out, using biopsy punches, and each disk was immersed in 50 μL of ultrapure water in a well of a 96-well PCR tray (BrandtTech®, Essex, CT, USA). For each filter membrane, 24 punches were examined, filling one quarter of a PCR-tray. The PCR-tray was then sealed and cooled down in an ethanol bath of a thermostat with a cooling rate of 1 K min^−1^, while a camera took pictures every 0.1 K from below. On these pictures, frozen wells can be well distinguished from unfrozen ones, and the cumulative number of frozen wells was assessed for the different samples, a clean filter and pure water. Concentrations of INP were calculated from the cumulative number of frozen droplets, based on the known amount of filtered water and Poisson statistics. A more thorough description of the measurement method and data evaluation was described earlier ^85^.

### Statistical analyses

Correlations between abundances of prokaryotic cells, small phototrophic eukaryotes and VLPs were investigated using R version 4.0.3. ^86^ within R studio^87^. Pearson or Spearman correlations were applied after the Shapiro-Wilk test confirmed normal distribution of data and residuals (for linear models). Dependences of EFs on environmental variables (wind speed, light, salinity) and interactive effects of those parameters were further investigated using linear regressions, and the models were validated using adjusted R^2^ and AIC in the R programming environment. Differences in alpha diversity and viral percent G/C base content were analyzed using a Kruskal-Wallis test with Dunn’s multiple comparison as *post hoc* analysis in Graphpad Prism v.9.4.1. Ecosystem-based differences in beta diversity shown in NMDS plots were assessed using PERMANOVA (n = 999 permutations) as well as Betadispersion analyses followed by a TukeyHSD test and executed by ‘adonis2’ and ‘betadisper’ function of the R package vegan ^88^, respectively.

### DNA extraction and sequencing of metagenomes

Genomic DNA was extracted from seawater (0.2 μm and <0.2 μm flocculated viral fraction), rain filter membranes (47 mm diameter, Merck/Sigma Aldrich, Munich, Germany), and the concentrated snow sample using the DNeasy PowerSoil Pro Kit (Qiagen, Hilden, Germany). DNA from aerosol filters (90 mm diameter, Merck/Sigma Aldrich, Munich, Germany) was extracted using DNeasy PowerMax Soil Kit (Qiagen) with a subsequent DNA precipitation step. After concentration in a speed-vac “Concentrator plus” (Eppendorf AG, Hamburg, Germany), DNA was quantified using Qubit™ dsDNA High Sensitivity Assay Kit on a Qubit™ 4 Fluorometer (Invitrogen/Thermo Fisher Scientific) and sent for metagenomic sequencing to Fulgent Genetics (CA, USA). Library preparation was done according to the Illumina DNA Prep with Enrichment Reference Guide (Document # 1000000048041 v05, June 2020).

### Metagenomic analyses

Raw shotgun sequencing reads of seawater (foams, SML, SSW), aerosols and precipitation datasets were quality-trimmed using bbduk (https://github.com/BioInfoTools/BBMap/blob/master/sh/bbduk.sh) and Sickle ^89^. Sequencing controls were assembled using MetaSPAdes version 3.13 ^90^ and used as a blueprint for read mapping ^91^ of actual samples; any reads that mapped to the negative controls were removed from downstream analyses (https://github.com/ProbstLab/viromics/tree/master/extract_unmapped_stringent). The same procedure including handling controls was carried out for metagenomic reads of aerosol samples.

Within a snakemake workflow ^92^ designed for detecting viruses and prokaryotes, quality-controlled paired-end reads were first assembled with MetaviralSPAdes ^93^ and reads were mapped back ^91^ to the assembly. Unassembled reads were assembled using MetaSPAdes version 3.14 ^90^, and the two assemblies were joined for downstream processing. VIBRANT v.1.2.1. ^94^, VirSorter v.1 ^95^ (only category 1, 2, 4, 5 were considered) and ViralVerify ^93^ were used to identify viral scaffolds and host contamination was removed with CheckV ^96^. Only viral scaffolds >10 kb were kept and clustered at species level (95 % similarity) using VIRIDIC ^97^, and the longest or circular scaffold of each cluster was used as representative. Metagenomic reads were mapped to >10 kb viral genomes with at least 90 % identity using Bowtie2 ^91^ with settings mentioned previously ^98^. To show the succession of two circular viral genomes across different samples, a separate mapping was done for these two scaffolds, and SNP analysis was performed for one marine virus that got airborne using Geneious v.11.1.5 ^99^ with default settings for variant analysis. Venn diagrams were constructed using ugent Webtool (https://bioinformatics.psb.ugent.be/webtools/Venn/). Complying to current viromics conventions ^100^, only scaffolds covered with 75 % of reads were considered further, and breadth was checked with Calcopo (https://github.com/ProbstLab/viromics/tree/master/calcopo). Mean coverage of viral scaffolds was calculated (https://github.com/ProbstLab/uBin-helperscripts/blob/master/bin/04_01calc_coverage_v3.rb), and sum-normalized based on sequencing depth. Genes on viral scaffolds were predicted using Prodigal v.2.6.3 ^101^ in meta mode and resulting coverages were further read-sum normalized and functionally annotated using DRAM-v ^102^. Synteny of viral genomes was visualized using Easyfig v.2.2.5 ^103^. Clustering of dereplicated viral genomes with a RefSeq database (release Dec. 2021) ^104^ was performed using vConTACT2 v.0.9.19 ^60,105^. Information on VCs and closest relative were compiled using graphanalyzer v.1.5.1 (https://github.com/lazzarigioele/graphanalyzer), and networks visualized in Cytoscape v.3.9 ^106^. Relative abundance of VCs was used for beta-diversity analysis. The % G/C base content of viral scaffolds counting towards a sample if detected based on read mapping, was calculated with an inhouse script (https://github.com/ProbstLab/uBin-helperscripts/blob/master/bin/04_02gc_count.rb). For investigating aerosolization and enrichment of VCs in rain over SML and foam, the maximum (sum-normalized) coverage of a virus across an ecosystem, e.g., across all aerosol samples, was considered assuming this value represents the highest possible abundance in that ecosystem. Then coverage ratios were calculated for the pairing rain/foam (R/F), rain/SML (R/S), aerosol/foam (A/F), aerosol/SML (A/S) and foam and SML samples were distinguished between the 5-0.2 μm and virome fraction.

### Prokaryotic community composition, binning of MAGs and virus-host interactions

Genes from combined scaffolds from the two assembly steps were predicted using Prodigal v.2.6.3 in meta mode ^101^, and genes were annotated using DIAMOND ^107^ blast against FunTaxDB ^58^. The genes were clustered at 99% identity using CD-HIT ^108^, and the centroid of the cluster was used for downstream mapping. Profiling of the prokaryotic community composition was done by read mapping using Bowtie2 of individual samples against scaffolds carrying *rpS3* genes. Taxonomic assignment of *rpS3* genes was performed using USEARCH ^109^ against the *rpS3* taxonomy database by Hug, et al. ^110^ Any unclassified and eukaryotic hits were excluded and coverages were read-sum normalized. For mean relative abundances taxonomic units of the same taxonomy were summed up. Analysis of the Shannon-Wiener Index (alpha diversity) using the ‘estimate_richness’ function, beta diversity, and NMDS plots based on Bray-Curtis dissimilarity were performed using phyloseq package^111^ in R version v.4.0.3.^86^ within R studio v.1.3.1093. Binning of MAGs was done using MetaBAT 2^112^ and Maxbin2^113^ with aggregating best genomes by DasTool ^114^ and followed by manual curation in uBin v.0.9.14. ^58^. MAGs were quality-checked using CheckM v.1.1.3 ^59^, and taxonomic assignment was performed with the classify workflow of GTDB-tk v.1.7.0 (database release r202)^115^. Mapping to individual MAGs was performed with Bowtie2 under allowance of 2 % error rate (3 mismatches) for breadth calculation. Virus-host interactions were inferred from CRISPR-spacer matches and shared k-mer frequency patterns between assembled viruses and host MAGs. At first, CRISPRCasFinder ^61^ with -minDR 16 was run on sample assemblies >1 kb and MAGs to find CRISPR consensus DR sequences from arrays with ≥ evidence level 3. Consensus DR sequences with 100 % similarity hits to viral scaffolds were removed from further analysis. Then DRs were used in MetaCRAST^116^ with settings -d 3 −l 60 -c 0.99 -a 0.99 -r to extract CRISPR spacer from the read files of each sample. Spacers were homopolymer and length-filtered (20-60 bp), clustered at 99% identity, BLAST was performed with a BLASTn –short algorithm ^117^ against the viral scaffolds, and filtered at 80% nucleotide similarity. Prokaryotic MAGs (116) were compared and dereplicated using dRep ^118^ at 95% average nucleotide identity. Additional MAGs that were excluded in the dRep process due to low quality in CheckM but had good contamination/completeness scores in uBin and formed their o– cluster in the dRep compare mode were additionally considered for k-mer based virus-host linkages. Viral scaffolds were assigned to these MAGs using VirusHostMatcher ^119^ at a d2* threshold of 0.3, as previously performed ^120^. Spacer-protospacer interactions and virus-host interactions based on k-mers were visualized using Cytoscape v.3.9 ^106^.

## Data availability

ECMWF ERA5 reanalysis data are freely available on https://cds.climate.copernicus.eu/cdsapp#!/home. Flow cytometry data have been stored at PANGAEA database and linked to the Integrated Marine Information System (IMIS). Sequencing data for this project are stored in Bioproject PRJNA811790, physical Biosamples SAMN26419398 – SAMN26419459 at NCBI. Sequence raw reads of samples and controls can be found at sequence read archive (SRA) under accession numbers SRR18243813 – SRR18243874. MAGs (116) are stored as Biosamples SAMN26866845 – SAMN26866960 and correspond to accessions JALHYV000000000 – JALHYZ000000000, JALHZA000000000 – JALHZZ000000000, JALIAA000000000 – JALIAZ000000000, JALIBA000000000 – JALIBZ000000000, JALICA000000000 – JALICZ000000000, and JALIDA000000000 – JALIDG000000000. The viral metagenome of 1813 scaffolds is stored within Biosample SAMN27124720, accession number JALJEC000000000. Data will be released upon publication of the manuscript. For further details, please refer to Table S10.

## Code availability statement

Links for relevant code are stated in the section “Online methods” or references for bioinformatic tools have been accordingly provided.

## Author contributions

JR conceptualized the study, conducted data analysis, wrote the first draft of the manuscript and together with SPE conducted field sampling. JP and AJP developed the viromics pipeline and together with SPE provided bioinformatic assistance. HAG measured and analyzed cell counts and together with MEH determined VLP counts, HW provided data on ice-nucleating particles in seawater samples. AS conducted binning of prokaryotic MAGs, CS and PG calculated backward trajectories. AJP provided supervision, bioinformatic guidance, help with analysis, scripts, and resources. All authors contributed to writing and editing of the final manuscript.

## Acknowledgements

We acknowledge funding for the MIDSEAS (Microbial dispersal from air to sea and snow) project by ASSEMBLE PLUS (European Union’s Horizon 2020 research and innovation program, Grant Agreement No. 730984), and by the German Aerospace Center (DLR) for the project DISPERS (50WB1922). JR received funding by the German Research Foundation (DFG RA3432/1-1). AJP received funding by the Ministerium für Kultur und Wissenschaft des Landes Nordrhein-Westfalen (“Nachwuchsgruppe Dr. Alexander Probst”) and the DFG (PR1603/2-1). MEH received funding within the framework of the PhD research-training group “The Ecology of Molecules” (EcoMol) funded by German Research Foundation within the TRR 51 Collaborative Research Center Roseobacter. HAG was funded the German Research Foundation (DFG), grant/award no. 34509606: Ökologie, Physiologie und Molekularbiologie der Roseobacter-Gruppe: Aufbruch zu einem systembiologischen Verständnis einer global wichtigen Gruppe mariner Bakterien. We like to thank the Sven Lovén Centre for Marine Sciences, Tjärnö, Sweden as part of the University of Gothenburg for hosting JR and SPE during the campaign and providing excellent science support including the use of laboratory and boat facilities. Here, we especially like to thank Anna-Karin Ring, Kerstin Johannesson, Joel White. We further thank Sabrina Eisfeld for laboratory maintenance, and Amelie Assenbaum for technical assistance during INP measurements. We also thank Ken Dreger for server administration, as well as Svenja Parge and Till Bornemann for bioinformatics assistance. The data handling was partially enabled by resources provided by the Swedish National Infrastructure for Computing (SNIC) at UPPMAX partially funded by the Swedish Research Council through grant agreement no. 2018-05973. Pavlin Mitev at UPPMAX is acknowledged for assistance.

