## Supplement material for "Heads in the clouds: marine viruses disperse bidirectionally along the natural water cycle"

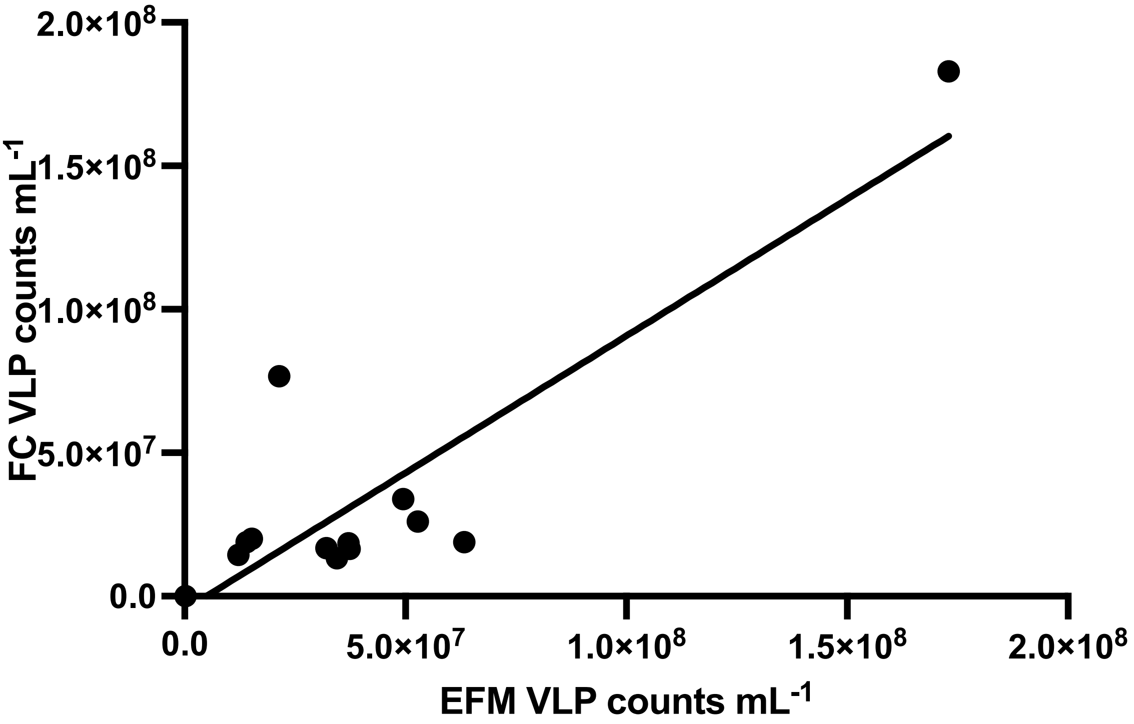


***Fig. S1: Linear regression for virus-like particles (VLP) from rain (n = 1), foam (n = 2), surface microlayer (n = 5) and subsurface water (n = 5) counted in flow cytometry (FC) versus virus-like particles counted under the epifluorescence microscope (EFM). The highest value corresponds to a foam sample.***

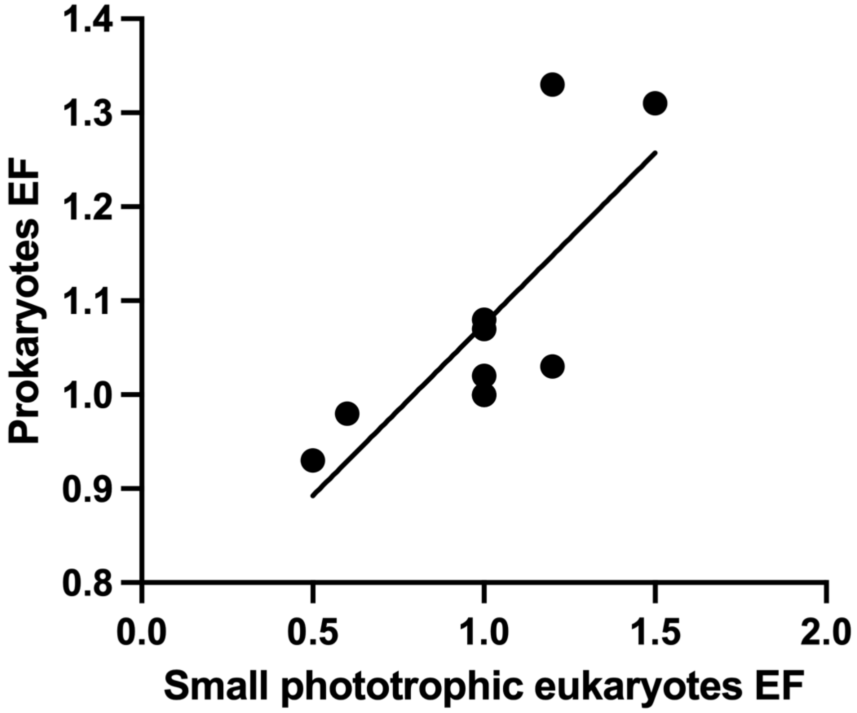


***Fig. S2: Enrichment factors (EF) of prokaryotic cells versus EF of small phototrophic eukaryotes in the surface microlayer over subsurface water.*** *Two-sided Spearman rank test indicates a significant (p = 0.0039, n = 10) correlation with Spearman's rho= 0.8264 between both parameters.*

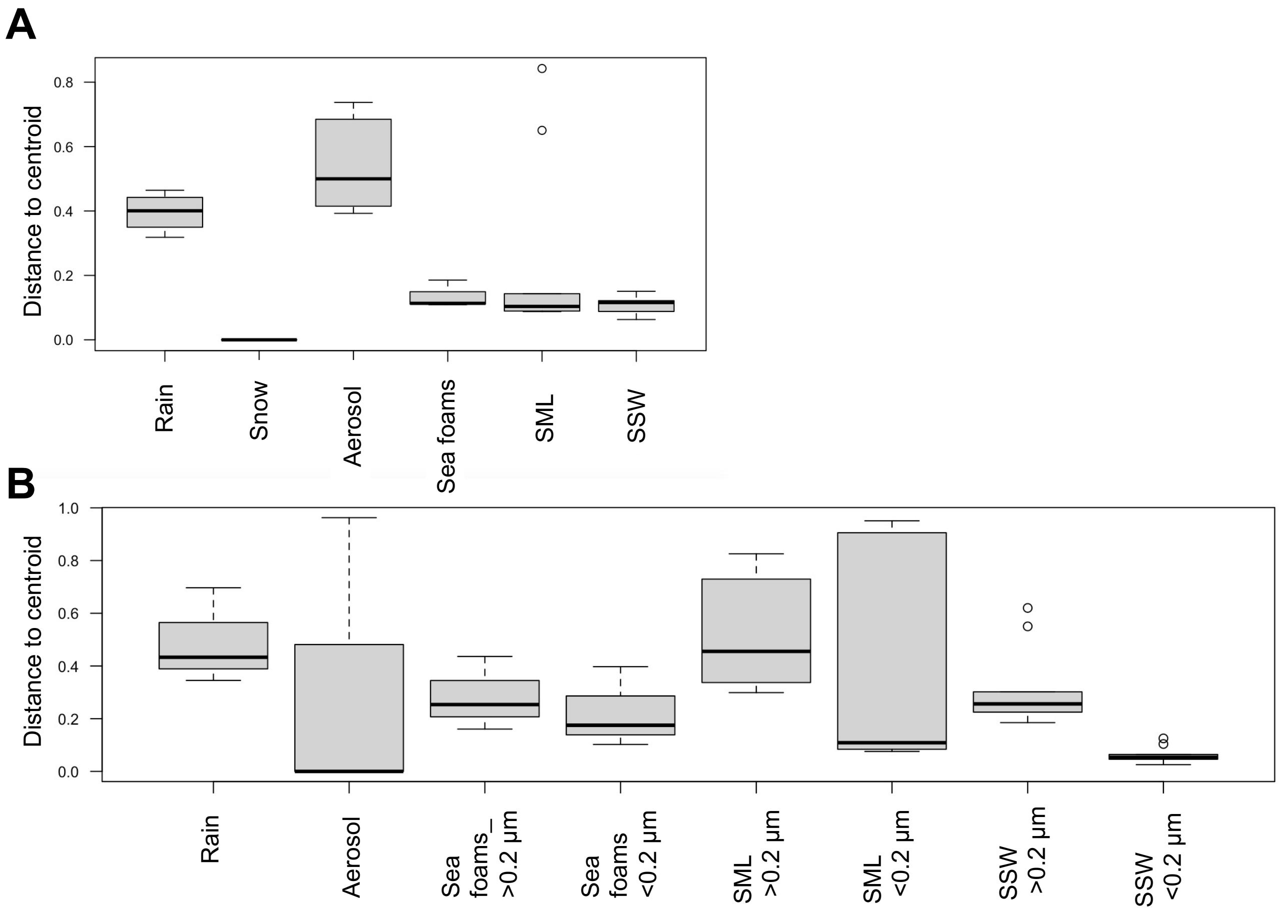


***Fig S3: Boxplots showing betadispers distances to centroid for various ecosystems and associated community of prokaryotes (A) and top 200 viruses (B).*** *Homogeneity of multivariate dispersions were calculated using the 'vegan' package in R and correspond to NMDS plots in Fig. 2B and Fig 3B. The center line represents the median, box limits are upper and lower quartile, whiskers indicate highest and lowest values and dots are outliers. SML=surface microlayer, SSW= subsurface water*

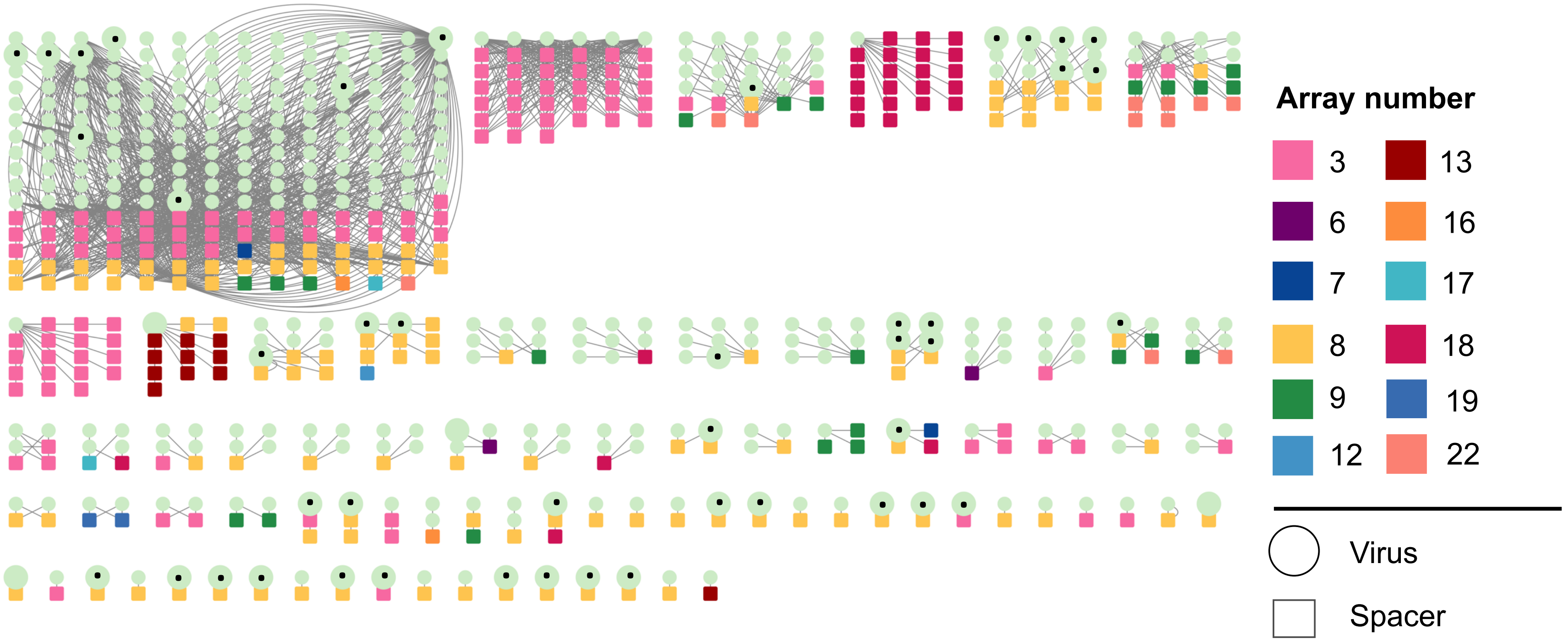


***Fig. S4: Matches of CRISPR spacers to viral protospacers separated by CRISPR array.*** *Squares represent spacers with different colors indicating different CRISPR array origins. Circles represent viruses with viruses assembled from rainwater being enlarged and unique rain viruses being further highlighted by a dot in the circle. Spacers from dominant arrays 3 and 8 match most viruses but also tend to match viruses from different ecosystems with array 3 and 8 providing spacers for marine and rainwater viruses, respectively.*

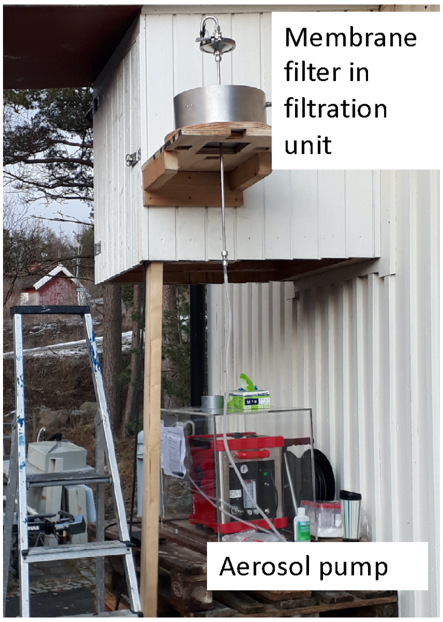


***Figure S5: Experimental set-up with aerosol pump and filter unit mounted at ~ 2 m above ground close to the coast.***

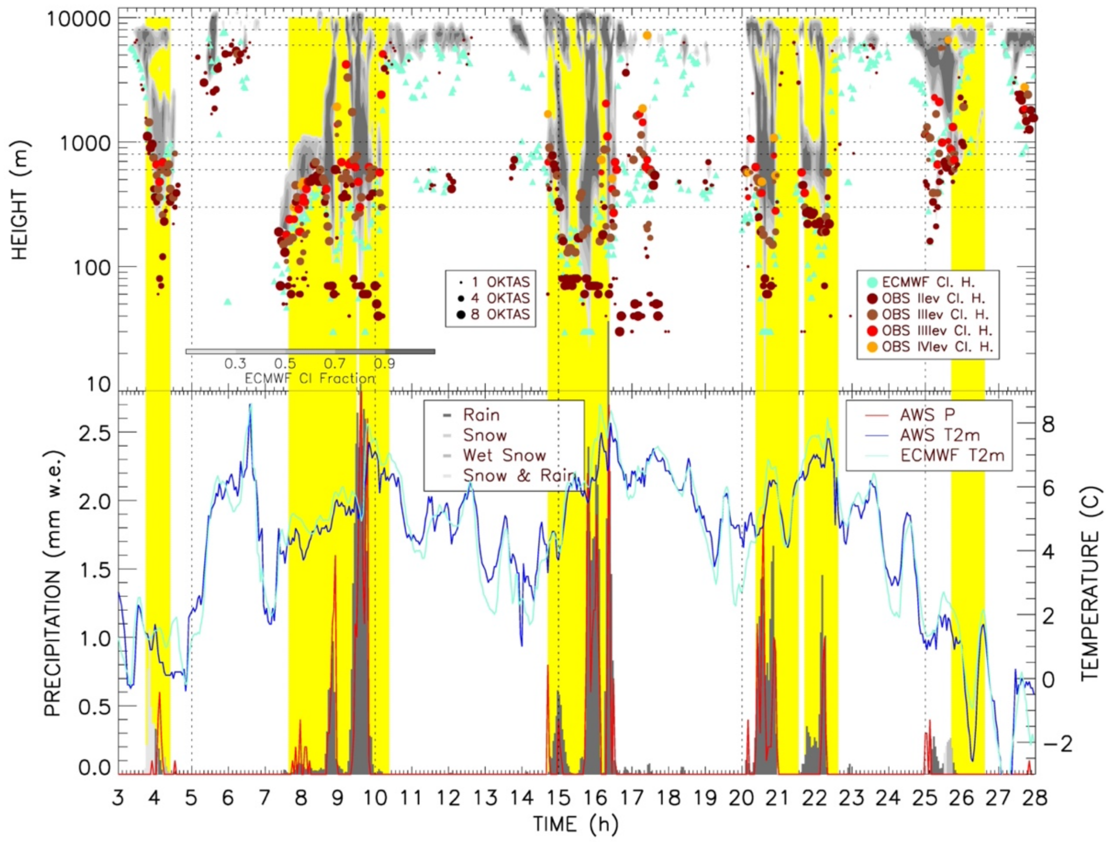


***Figure S6: Meteorological and cloud conditions at Nordkoster A Automatic Weather Station (AWS) during February 2020.*** *Upper panel ECMWF ERA5 cloud fraction (filled contour), and cloud base height from model (sky blue filled triangles) and from in situ observations (colored filled circle). In situ cloud base observations are divided on the base of altitude level (color of filled circle) and cloud coverage at that level (size of the filled circle). Lower panel shows 2-meter temperature from AWS (blue line) and ECMWF ERA 5 (sky blue line). Vertical stacked bars in different gray scale color highlight quantity and phase of ECMWF ERA 5 precipitation field whereas red line show precipitation measured by AWS. Rain sampling intervals are highlighted with yellow areas.*

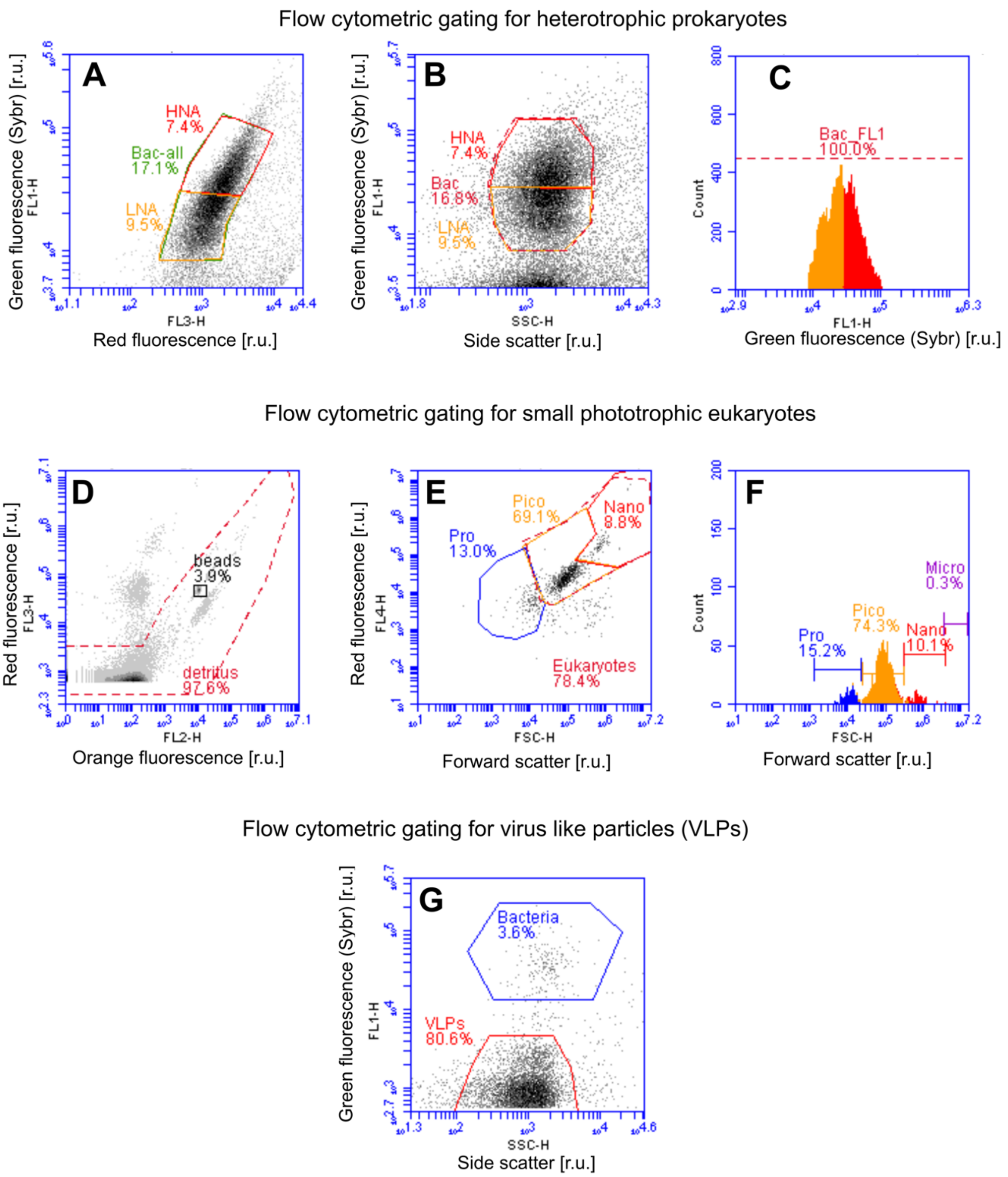


***Fig. S7: Flow cytometric gating strategies for heterotrophic prokaryotes (A-C), small phototrophic eukaryotes (D-F) and virus-like particles (VLPs, G) using a BD Accuri C6 flow cytometer (Becton & Dickinson Biosciences) with its Flow C software.*** *Discrimination of phototrophic prokaryotes (A), double-check of (sub-)populations against forward scatter and their distribution as histogram (C). Discrimination of detritus, internal beads, and background noise for the autofluorescence of small phototrophic eukaryotes (D), re-gating of all events except detritus (E) and histogram of the subpopulations of phototrophic pico-, nano- and microplankton (F). Phototrophic prokaryotes were discriminated from the analysis due to the insufficient detection by the Accuri C6 after Ribeiro et al. (2016). r.u., relative units.*

***References:***

Ribeiro, C.G., Marie, D., dos Santos, A.L., Bandini, F.P., and Vaulot, D. Estimating microbial populations by flow cytometry: Comparison between instruments. *Limnol. Oceanogr. Methods* **14**, 750-758. doi: 10.1002/lom3.10135 (2016).
