## Extended Data for "Heads in the clouds: marine viruses disperse bidirectionally along the natural water cycle"

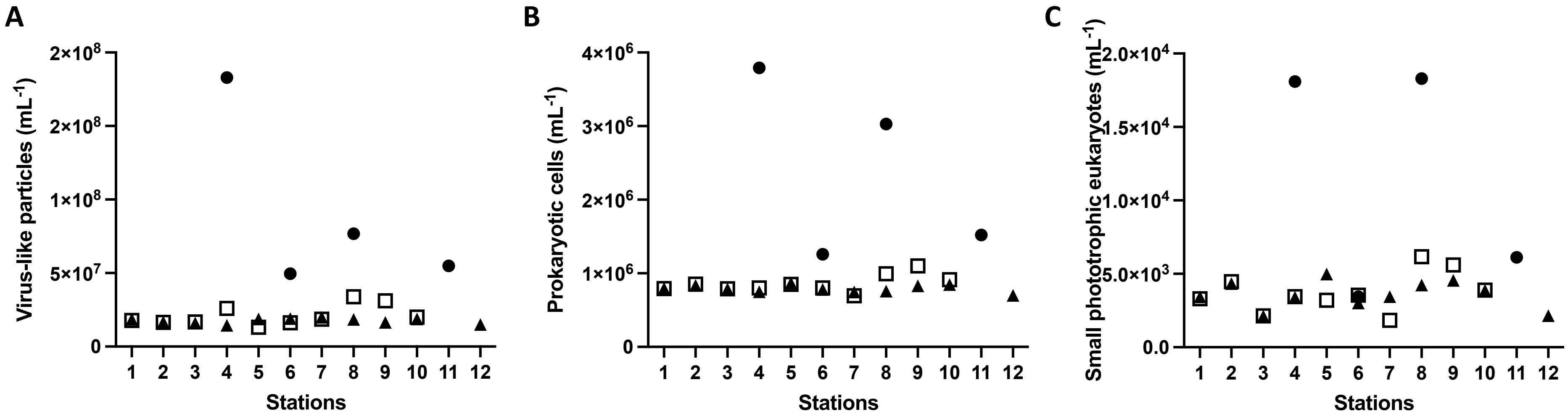


***Extended Data Fig. 1: Absolute counts of virus-like particles (VLP) A), prokaryotic cells B) and small phototrophic eukaryotes mL$^{-1}$ C) as counted in the flow cytometer across 12 sampled stations. SML= surface microlayer, SSW= subsurface water***

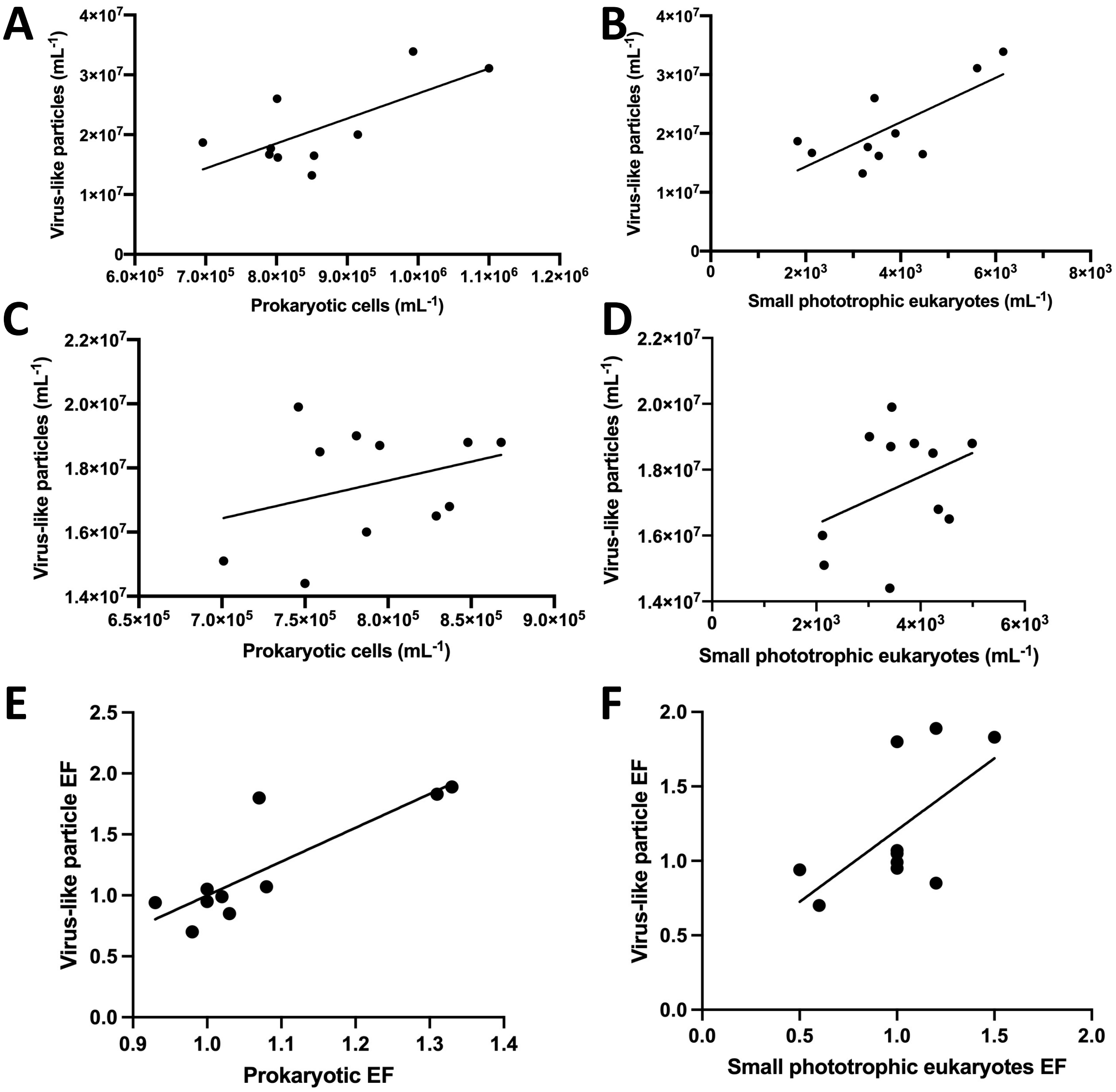


***Extended Data Fig. 2: Relationship of virus-like particles (VLP) and host cells in the neuston and the plankton.*** *Linear regression for VLP versus prokaryotic cells (A) and small phototrophic eukaryotes (B) in the surface microlayer corresponding to the neuston. Linear regression for VLPs versus prokaryotic cells (C) and small phototrophic eukaryotes (D) in the subsurface water corresponding to the plankton. Linear regression for VLPs enrichment factors (EF) versus prokaryotic EF (E) and small phototrophic eukaryotes EF (F) for the surface microlayer compared to subsurface water.*

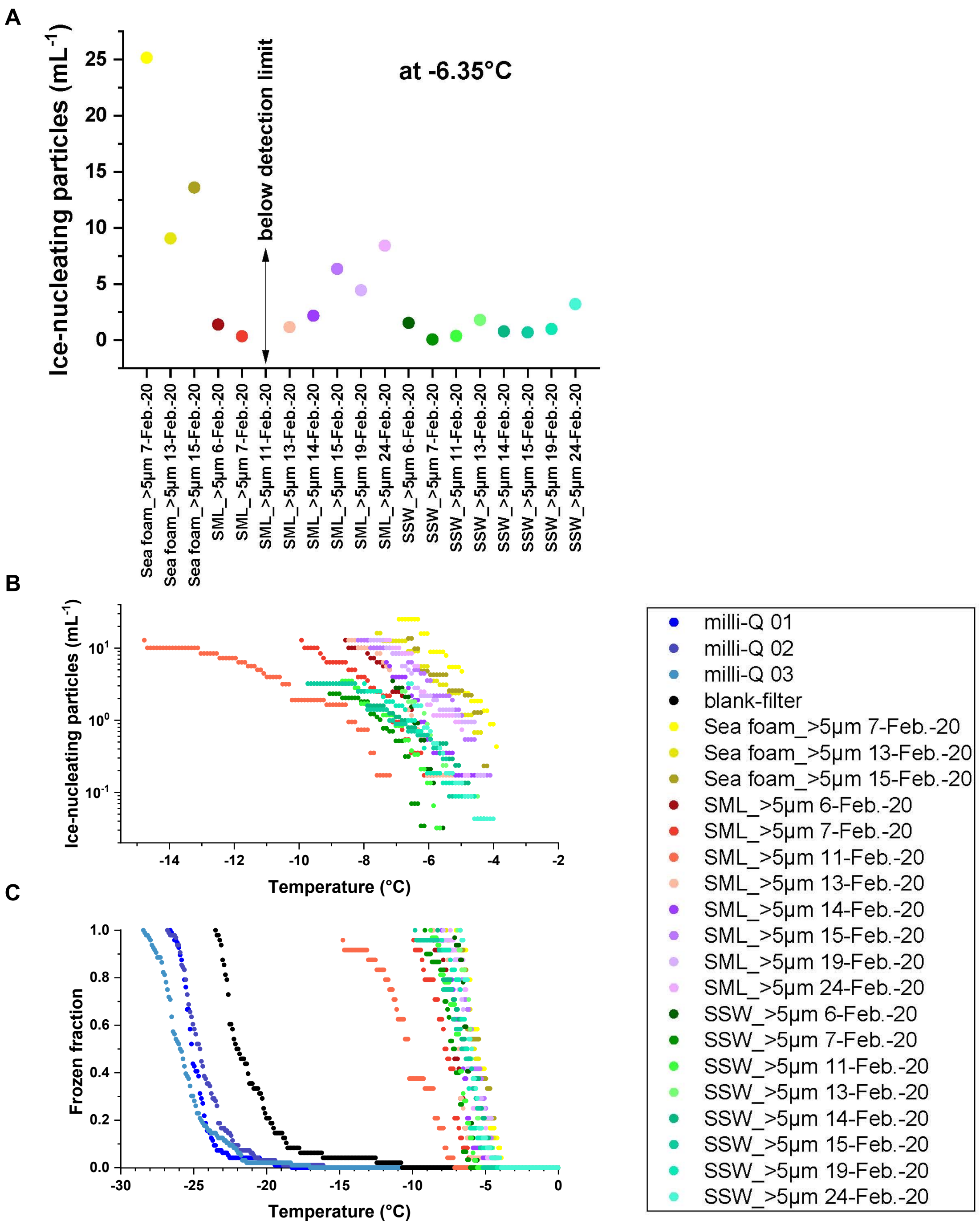


***Extended Data Fig. 3: Results from measurements of ice-nucleating particles (INP) in foams, surface microlayer (SML) and subsurface water (SSW).*** *INP concentrations at the ice nucleation temperature of -6.35°C (A), INP spectra over the detectable temperature range (B) and the measured frozen fraction values in comparison with results from pure water (C) showing that all samples were well above the background.*

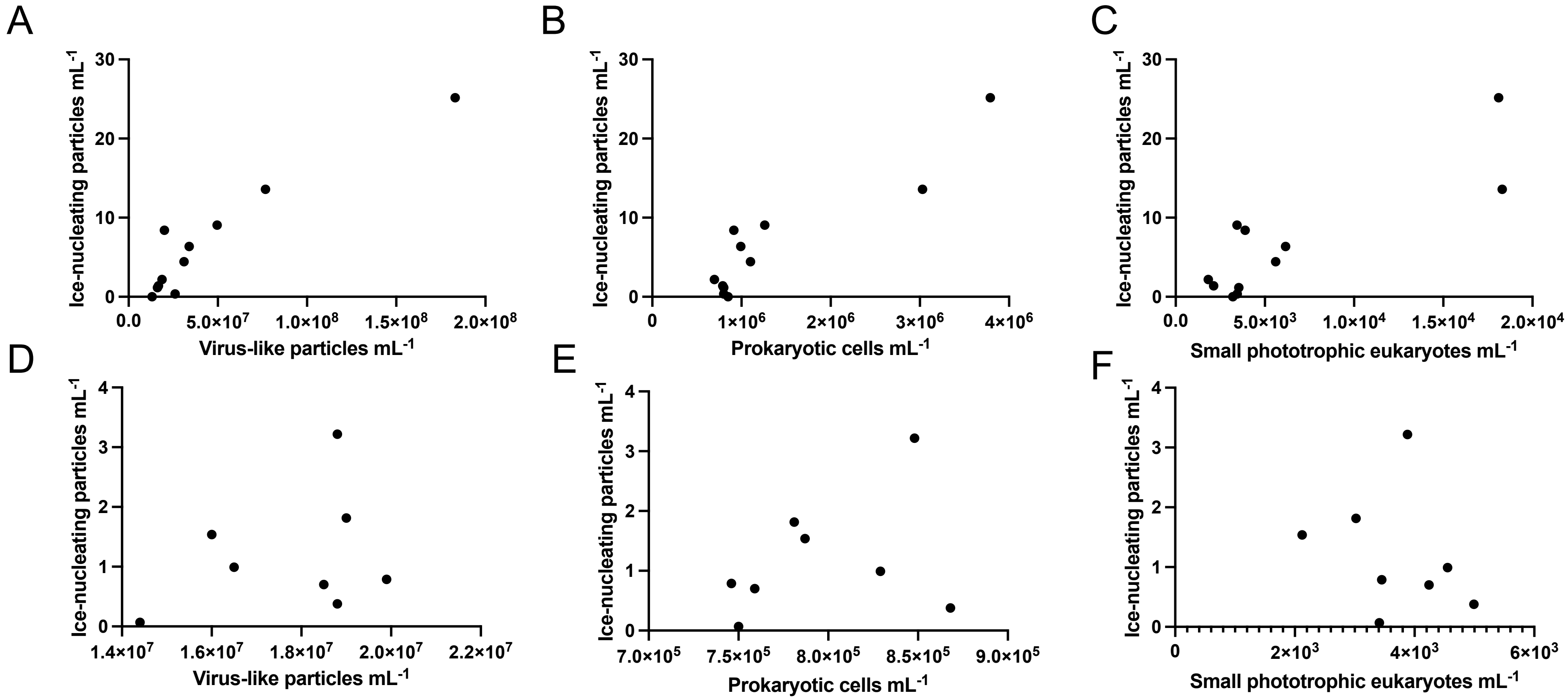


***Extended Data Fig. 4: Ice-nucleating particles (INP) in foams, surface microlayer (SML) and subsurface water (SSW) and their relationship to cell and virus-like particle (VLP) numbers.*** *Correlation of INP versus VLP, prokaryotic cells and small phototrophic eukaryotes in the neuston (A-C) and in the plankton (D-F). Data were not normally distributed for A-C but for D-F. Spearman rank correlation coefficients and p-values were 0.86 and 0.0012, 0.79 and 0.0055, 0.65 and 0.037 for A-C, respectively and Pearson correlations were not significant ($p > 0.05$) for D-F.*

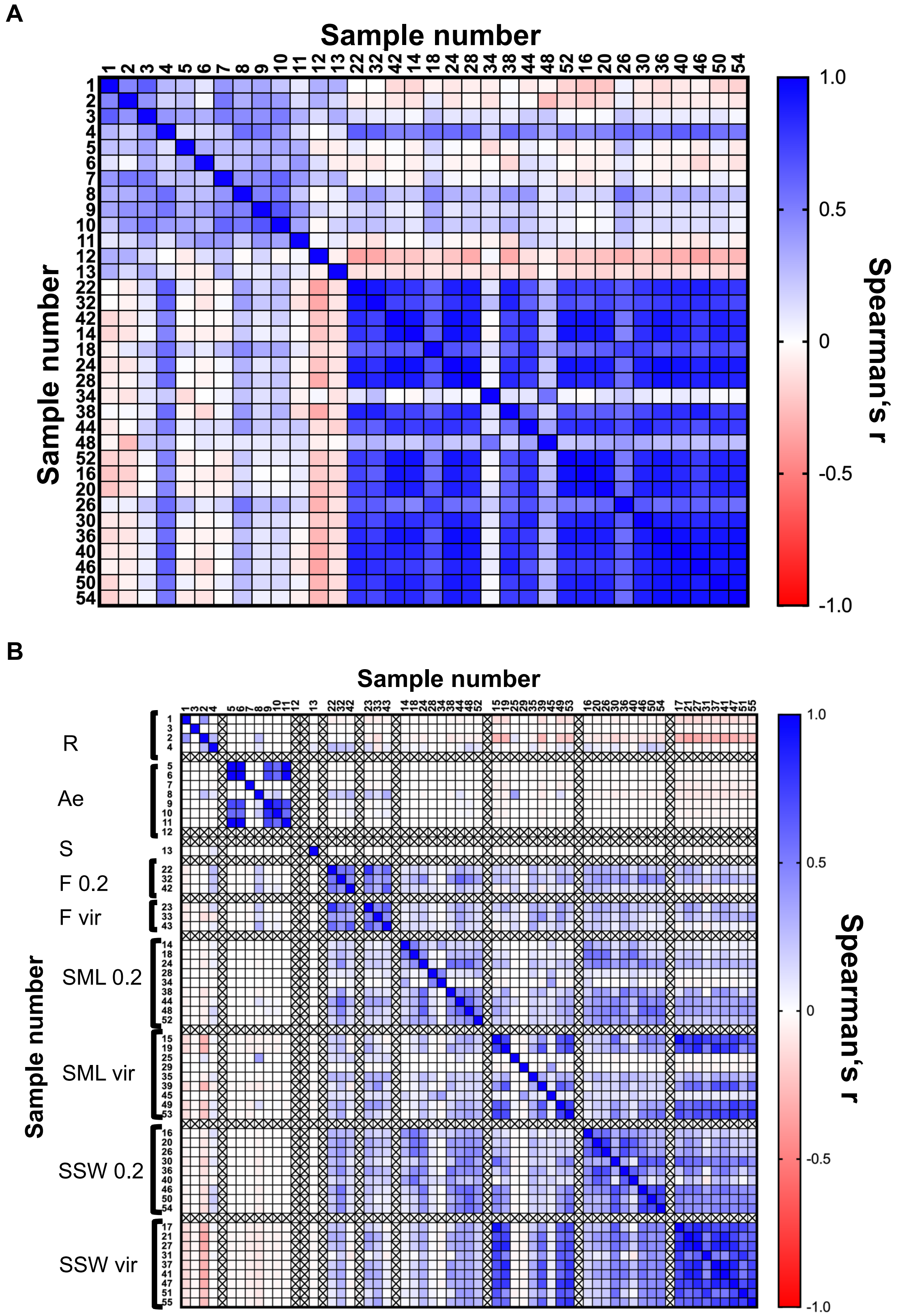


***Extended Data Fig. 5: Correlation matrix for relative abundances (based on read-normalized coverage) of prokaryotic taxa (n=69) detected by rps3 gene prediction as shown in Figure 3 across rain, snow, aerosol, foam, surface microlayer (SML) and subsurface water (SSW) samples***. *(A). Correlation matrix for abundances (based on read-sum normalized coverage) of 1813 viral scaffolds across rain (R), snow (S), aerosol (Ae), foam (F), SML, and SSW samples*

*(B). Sample numbers are in accordance with Table S11, vir = viromes, 0.2 = 0.2 µm fraction. Spearman rho, p values and confidence intervals for each pairwise comparison (two-sided test) shown in the correlation matrices are given in Appendix 1 and 2 of the Supplement Tables.*

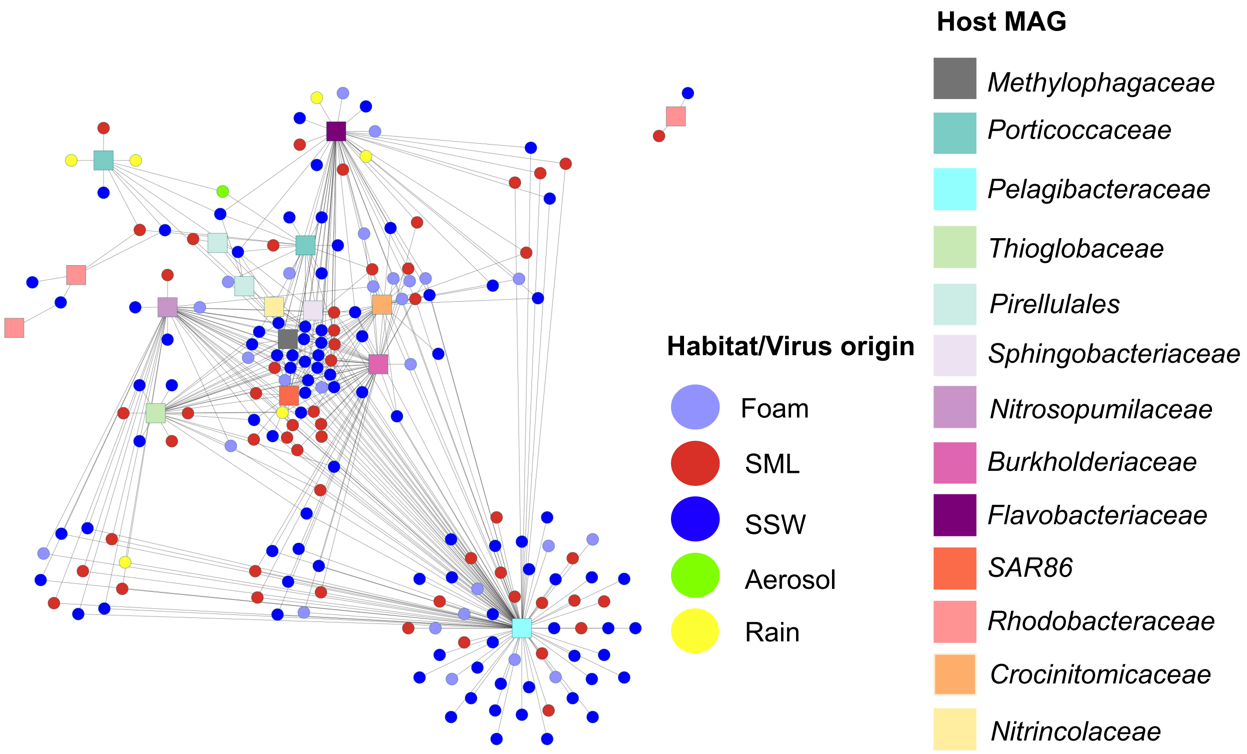


***Extended Data Fig. 6: Virus-host interaction based on k-mer frequency patterns.*** *Using VirhostMatcher at a d2* dissimilarity threshold of < 0.3, viral scaffolds (represented as circles) were tested against a set of 26 dereplicated host MAGs (squares). Visualization was performed in Cytoscape v.9.3.*

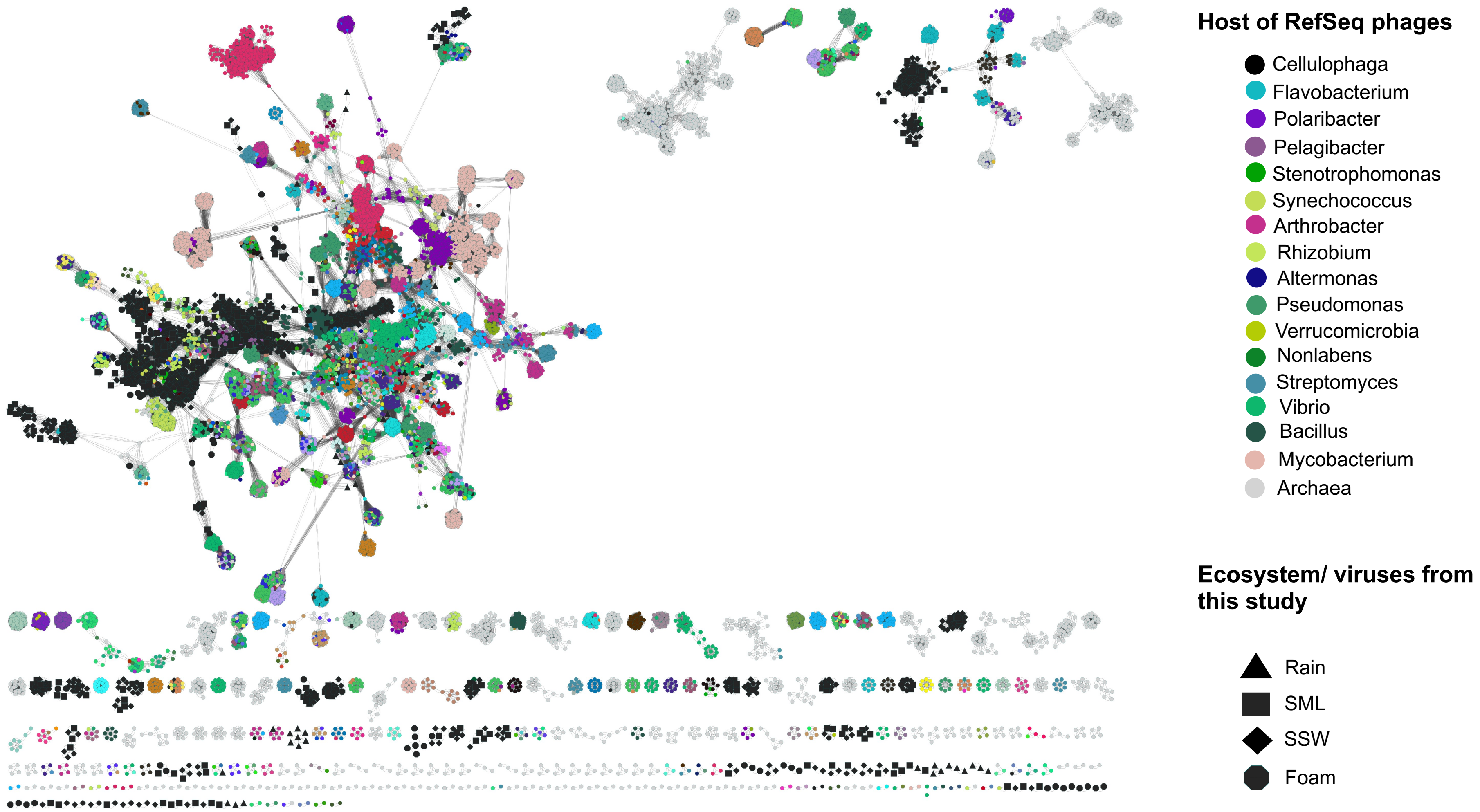


*Extended Data Fig. 7: Viral clustering of viruses from different ecosystems with Refseq database (December 2021) reveal many clusters with unrelatedness to Refseq db. The legend has been reduced to hosts that show interactions with viruses from this study.*
